# Multimodal Approach for Identification and Validation of Hepatocellular Carcinoma Targets for Radiotheranostics

**DOI:** 10.1101/2025.11.21.689799

**Authors:** Philip Homan, Joon-Yong Chung, Woonghee Lee, Divya Nambiar, Julia Sheehan-Klenk, Hima Makala, Stanley Fayn, Orit Jacobson, Arthur Paden King, Kyungeun Kim, Jeong Won Kim, Sung Ryol Lee, Maggie Cam, Stephen M. Hewitt, Xin Wei Wang, Mitchell Ho, Freddy E. Escorcia

**Author notes:** Corresponding author. Email: Freddy E. Escorcia, MD, Ph.D. These authors contributed equally to this work (Philip Homan, Joon-Yong Chung).

## Abstract

Identifying tumor selective targets is critical for the development of precision diagnostic and therapeutic agents in oncology. Despite advances in precision oncology elsewhere, there are no FDA-approved hepatocellular carcinoma (HCC)-selective treatments. HCC is the most common type of liver cancer and accounts for significant morbidity and mortality worldwide. Here, we sought to integrate bulk (371 cases) and single cell RNA sequencing (scRNAseq, *n*=2 datasets, 34 cases, 102,956 cells) of patient samples to enrich for molecules that are overexpressed in HCC, which could serve as HCC-selective targets. To guide definitions of tumor and normal cell clusters with higher fidelity, we also imported a normal liver scRNAseq dataset. Using this integrated approach, we identified several HCC-selective plasma membrane molecules. To validate these targets, we performed immunohistochemical staining of HCC and normal tissue microarrays and confirmed HCC-selective staining of identified targets. Next, we verified the presence of these targets in several commercially available HCC cell lines by flow cytometry and western blot. Finally, we designed, engineered, and tested novel antibody-based positron emission tomography (immunoPET) agents to these targets in various murine models of liver cancer. Our findings confirm that we can leverage this multimodal approach to identify and validate of HCC-selective targets, which can be used to develop tumor-selective diagnostic and therapeutic radiopharmaceuticals, or radiotheranostics, and other precision oncology agents.

**One Sentence Summary:** A multimodal pipeline defines and validates tumor-selective surface targets for radiotheranostic use in hepatocellular carcinoma.

## INTRODUCTION

Hepatocellular carcinoma is the most common cause of liver cancer and accounts for over 500,000 new cases and as many deaths annually worldwide (*1*). While transplantation and surgical resection can be curative, and several locoregional therapies are available for early-stage disease, most patients present with advanced disease or metastatic disease. For these patients, systemic agents are indicated (*2*).

Despite advances in precision oncology, there are no clinically approved HCC-selective treatments. Targeted therapies in the form of kinase inhibitors were the first to significantly improve survival for patients with HCC (*3–5*). More recently, immunotherapy-based regimens have come to the forefront, demonstrating superior outcomes compared to kinase inhibition alone (*6, 7*). While these agents have fundamentally changed our approach in managing patients with advanced and metastatic HCC, the 5-year overall survival of this cohort is still only about 20 percent. Furthermore, this class of agents is not truly precision oncology, where a unique feature, such as a novel oncogenic fusion or overexpressed molecule, of the cancer is specifically exploited to yield superior therapeutic benefit—that is, improved overall survival and/or quality of life for patients receiving the novel therapy.

In other malignancies antibody-drug conjugates (ADCs) targeting HER2, TROP2, and Nectin-4 have been successfully tested in clinical studies, yielding some regulatory approvals in breast (*8, 9*), and urothelial cancers (*10*), respectively. Additionally, targeted antibodies specific to CD20, peptides specific to somatostatin receptors (SSTR), and peptidomimetics specific for prostate specific membrane antigen (PSMA), and small molecules targeting norepinephrine receptors have been coupled to therapeutic radioisotopes are clinically approved for patients with recurrent B-cell lymphomas (*11–13*), gastroenteropancreatic neuroendocrine tumors (*14*), metastatic castration resistance prostate cancers (*15*), and paragangliomas (*16*), respectively.

Some therapeutic targets, including Glypican-3 (GPC3) and basigin (BSG, CD147), plasma membrane molecules known to be overexpressed in HCC, have been explored in preclinical and clinical trials (*17–21*). Imaging of GPC3 using an antibody-based positron emission tomography agent (immunoPET) proved successful in a clinical trial of patients with HCC, however, the imaging agent was found to not predict for response to kinase inhibition (*22*). Preclinically, Fu et al. engineered a GPC3-targeted ADC and demonstrated efficacy in murine models (*23*). Others have successfully used alpha particle-emitting radiopharmaceutical therapy to treat animals with GPC3-expressing liver cancer xenografts (*24–26*). Early clinical data using a peptide-based positron-emission tomography agent appear promising (*27*). In China, a radioiodine-labeled CD147-specific antibody fragment, ^131^I-metuximab, is approved for transarterial adjuvant treatment following resection for patients with HCC (*21*). Because HCC is radiosensitive as it is routinely treated with external beam radiotherapy and transarterial radioembolization, it stands to reason that targeted radiopharmaceutical therapy may represent a viable treatment option for patients with this disease. The successes targeting HCC-selective molecules for therapeutic intent suggest that an approach to identify additional targets could yield novel therapeutics and diagnostics, also referred to by the portmanteau “theranostics” or “radiotheranostics” when specifically using radionuclides, for patients (*28, 29*).

Tumor plasma membrane associated targets are of particular interest for radiotheranostic development because they are easily accessible to parenterally administered agents. To the best of our knowledge, no systematic evaluation of plasma membrane targets has been performed to generate a library of putative targets for HCC. Such an approach may help inform combinatorial treatments and begin to address the known challenges of tumor heterogeneity in the development of therapeutic agents.

Here, we perform *in silico* screens of bulk (The Cancer Genome Atlas, TCGA) and single cell RNA sequencing (scRNAseq) patient datasets to identify a list of plasma membrane molecules that are overexpressed on HCC compared to non-tumor samples. We then assess whether these putative targets are also present on patient tissue microarrays (TMAs) of tumors and normal tissues, and evaluate their expression in commercially available liver cancer cell lines by flow cytometry and western blot. Next, we design, synthesize, and successfully test immunoPET agents to select targets and confirm their utility for imaging in several murine models of liver cancer. Together, this work demonstrates proof-of-concept that our target discovery approach can yield tumor-selective agents that could be engineered for radiotheranostics and other diagnostic or therapeutic applications.

## RESULTS

### Bulk and single cell RNAseq analysis from patients with HCC identify differentially expressed plasma membrane-associated genes

To identify potential plasma membrane genes associated with liver hepatocellular carcinoma, we compared the gene expression profiles of LIHC tumor samples from TCGA to normal liver tissue. The dataset consisted of 371 tumor and 50 normal liver samples. PCA analysis of the tumor showed that the tumor and normal liver samples did differentiate into separate clusters (**Fig. S1A**). Differential Expression analysis between Tumor and Normal samples was performed on cell surface associated marker genes, as defined by the UniProt database. The analysis identified 156 genes differentially expressed in the tumor samples (**Fig. 1A**).

**Fig. 1.**
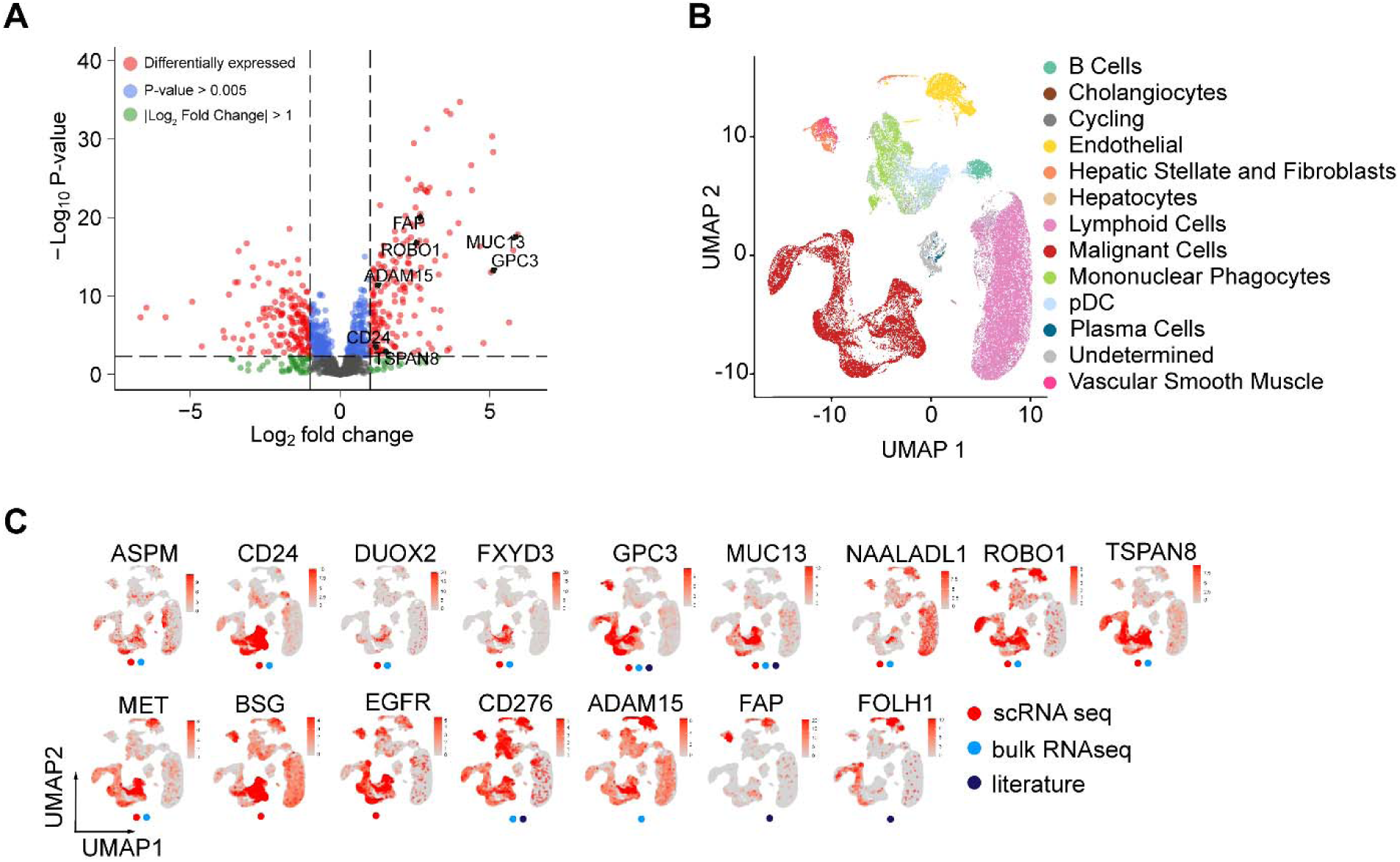
Identification of overexpressed Genes in HCC. **(A)** Volcano plot of differentially expressed genes from TCGA-LIHC RNA-seq expression data. Genes colored in red are identified as differentially expressed (*p* value <0.005 and |Log_2_ Fold Change| >1). Genes colored in green have a |Log_2_ Fold Change| >1 and genes colored in blue have P-value <.005. *MUC13*, *GPC3*, *CD24*, *TSPAN8*, *ADAM15*, *ROBO1*, *FAP* are highlighted. **(B)** UMAP of all cells from 41 HCC tumor samples and 8 normal liver samples. Cells are colored by major cell type determined from the Human Single Cell Atlas. Malignant Cells (Red) were identified from the Lichun Ma *et al.* (35) dataset. **(C)** UMAP plots showing the expression levels of select genes. Plots are marked by red dot if the gene was identified as differentially expressed in the single Cell RNA-seq dataset, blue differentially expressed in RNA-seq dataset, or dark blue if identified in the literature.

To help identify high quality markers directly associated with tumor malignancy, we examined previously published single cell datasets associated with hepatocellular carcinoma. 31 HCC tumor samples published in Lichun Ma *et al.* (*30*) were integrated with 10 HCC tumor samples and 8 normal liver samples published in Yiming Lu *et al.* (*31*). After quality control filtering of each sample the integrated dataset contained 102,956 cells. We performed unsupervised clustering and annotated cells using the data from the Liver Single Cell Atlas (*32*), which contained 28 human liver biopsies from 5 different studies (**Fig. 1B**). After integration, we found that the relative cell type proportion remained consistent between both datasets (**Fig. S1B**). Malignant cells were previously identified within tumor liver samples by inferring large scale chromosomal copy-number variations (CNVs) (*30*). All malignant cells from this study clustered into a distinct population. Twenty-five percent of the tumor cells from the Yiming Lu *et al.* dataset also clustered within this population and were annotated as malignant. To confirm the identity of the malignant cells, normal liver samples from this dataset were included. Less than 10% of the normal tissue cells clustered with the malignant cell population (**Figs. S1C, 1D**).

We then compared the malignant cells to all other liver cell populations to identify genes specifically expressed in malignant cells. Genes related to cell surface markers as identified by the UniProt (*33*) database were selected from the differentially expressed genes. 63 differentially expressed cell surface markers specific to the malignant cell population were identified. An overlay of gene expression from select differentially expressed genes extracted from the current analysis and literature shows that expression of some is enriched in malignant cells, though others had broader expression in normal cells (**Fig. 1C**).

We found that 10 genes were classified as both malignant cell-specific in the single cell HCC samples and tumor specific from the bulk RNAseq dataset of TCGA (**Fig. S2A**). These genes show a high specificity for the malignant cell population (**Figs. 1C, S2B**). CD24, GPC3, and TSPAN8 each target approximately 50 percent of malignant cells.

### Gene expression from Human Protein Atlas helps Identify liver specific genes

We further investigated the possibility of non-liver tissues expressing targets identified in the malignant cell differential genes. We collected gene expression data from the Human Protein Atlas database (PADB) and calculated the fold change between the gene expression from 51 human tissues compared to the expression in liver tissue. Genes with a low score are considered liver specific and high-quality targets (**Fig. S2C**). This metric provides a simple insight into the specificity of identified genes for HCC tumor cells. ASPM, ROBO1, GPC3, and MUC13 were confirmed as good tumor liver targets while CD24, SLC6A8, and FXYD3 showed extensive off tissue expression.

### *In silico*-identified theranostic markers are highly expressed in HCC patient tissue microarrays and associate with clinical outcomes

Immunohistochemistry was performed on hepatocellular carcinoma (HCC) tissue microarrays to assess the clinical significance of eight candidate therapeutic markers. Seven targets (CD24, GPC3, MUC13, TSPAN8, MET, CD147, and EGFR) were identified through *in silico* analysis, and ROBO1 was included based on a prior literature report (*17*). Expression levels were evaluated based on their localization to either the membrane or cytoplasm, using digital image analysis (**Fig. 2A**). Among these markers, CD24, GPC3, and MUC13 demonstrated high membrane expression in 24.9% (41/165), 63.0% (104/165), and 52.7% (87/165) of HCC samples, respectively (all *p*<0.0001), but were undetectable in adjacent non-tumor liver tissues (**Figs. 2B, 2C**). Similarly, TSPAN8, MET, and EGFR were predominantly expressed on the membranes of tumor cells in 33.3% (55/165), 77.6% (128/165), and 61.2% (101/165) of cases, respectively, with minimal presence in non-tumor regions (all *p*<0.0001; **Figs. 2B, 2C**). ROBO1 and CD147 also showed significantly elevated membrane expression in HCC tissues (both *p*<0.0001), but their expression in non-tumor liver tissues was comparable, suggesting baseline expression of these markers in normal hepatocytes (**Figs. 2B, 2C**).

**Fig. 2.**
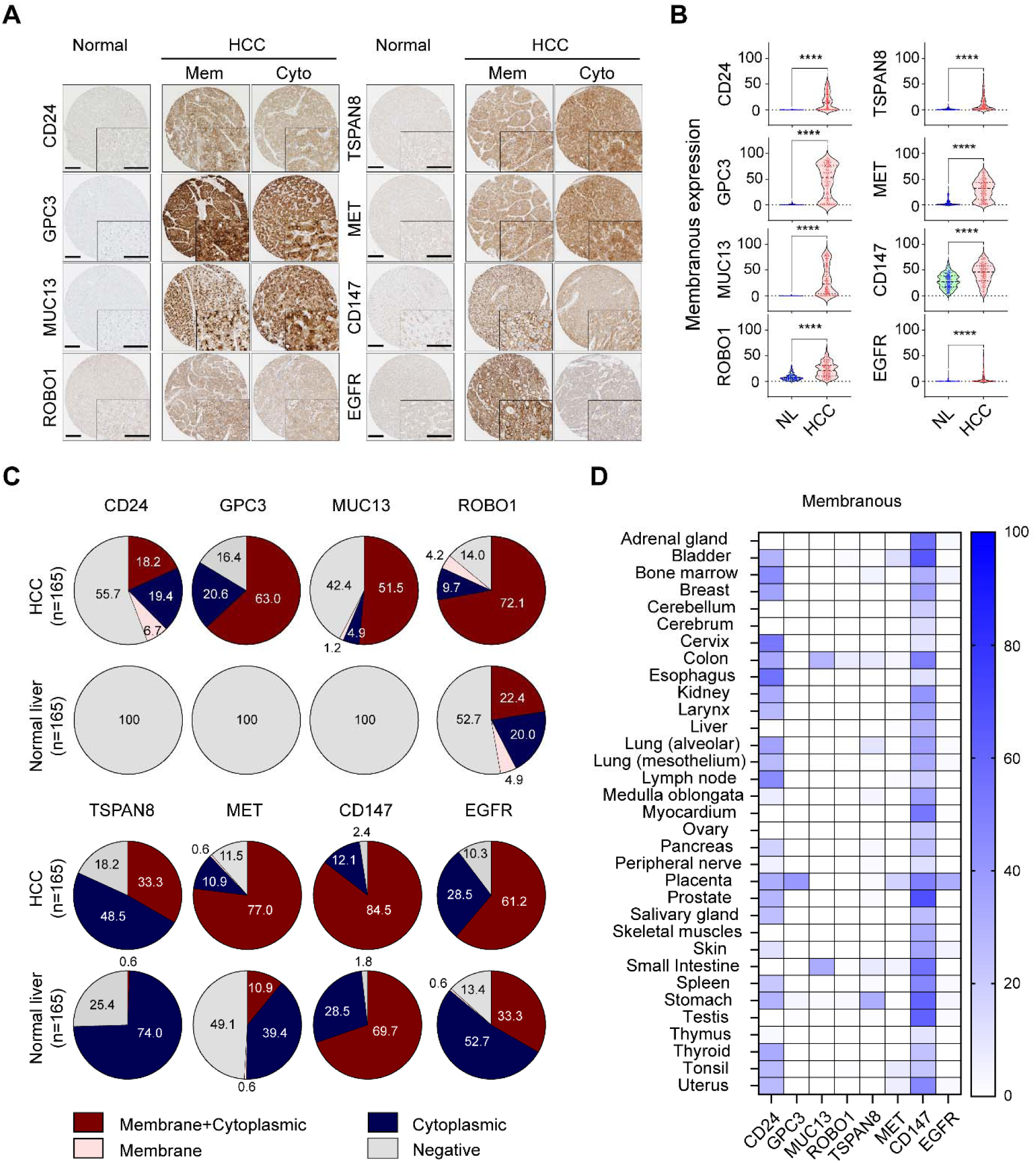
Profiling of potential theranostic markers in human HCC and normal tissues. (**A**) Representative immunohistochemical images showing the expression of CD24, GPC3, MUC13, ROBO1, TSPAN8, MET, CD147, and EGFR in formalin-fixed, paraffin-embedded sections of normal and HCC tissues. High magnification images are presented in the inset. Mem, membrane; Cyto, cytoplasm. Scale bar = 100 *µ*m. (**B**) Violin plots depicting membranous expression levels of eight potential theranostic markers in 165 HCC patient samples and corresponding non-adjacent normal tissues. Statistical significance was assessed by two-tailed t-test. ****, *p*<0.0001. (**C**) Pie chart illustrates the distribution of expression patterns for the eight theranostic targets, categorized as combined membrane and cytoplasmic, cytoplasmic, membrane, and negative. Data are shown for both HCC tissues (upper panel) and matched non-adjacent normal tissues (lower panel). (**D**) Heatmap comparing protein expression in normal human tissues based on membrane localization. The heatmap reflects relative protein expression, with darker blue indicating higher levels and lighter blue representing lower levels.

Next, we examined the correlation between the marker expression and patient survival outcomes. Kaplan-Meier survival analyses demonstrated that patients with high CD24 (*p*=0.025), MET (*p*=0.023), and CD147 (*p*=0.005) membranous expression, as well as those with low MUC13 (*p*<0.001) membrane expression, had significantly shorter DFS (**Fig. S3A**). Additionally, high TSPAN8 expression correlated with poorer OS (*p*=0.045; **Fig. S3B**). Cox proportional hazards analyses further confirmed that high MET (HR=1.727; 95% CI: 1.076–2.773; *p*=0.024) and high CD147 (HR=1.869; 95% CI: 1.199–2.912; *p*=0.006) expression were significantly associated with worse DFS in univariate analysis. Conversely, high MUC13 expression (HR=0.386; 95% CI: 0.232–0.644; *p*<0.001) correlated with improved DFS (**Table S1**).

Furthermore, pathologic T-stage (pT) and distant metastasis were identified as independent prognostic factors for DFS, while pT stage alone remained an independent prognostic indicator for OS (**Table S1**). Cytoplasmic expression was also analyzed and is summarized in **Figures S4** and **S5**. Importantly, while expression-survival correlations can be hypothesis generating, further validation is needed.

In the selection of tumor-selective targets for therapeutic development, it is critical to assess protein levels in normal tissue as well to rule in or out potential (e.g. on target, off tumor) toxicity. Accordingly, we assessed their presence across various normal human tissues using normal tissue microarrays. Immunohistochemical analysis quantified both membranous and cytoplasmic expression. Membranous GPC3 and EGFR were predominantly expressed in the placenta, with minimal detection in other normal organs (**Fig. 2D**). Notably, MUC13 and TSPAN8 showed peak expression in the colon and small intestine, respectively, while ROBO1 had negligible expression across all tested normal organs. Moderate MET expression was observed in the placenta and bladder. Meanwhile, CD24 and CD147 displayed moderate to strong membranous expression across multiple tissues. In sum, our normal tissue TMA assessment suggests that GPC3, MUC13, ROBO1, TSPAN8, MET and EGFR appear to exhibit low normal tissue expression at the protein level, suggesting that these represent promising theranostics targets for HCC. However, CD24 and CD147 may be less attractive for such purposes, unless alternate modes of administration (e.g., transarterial) are considered to avoid off-tumor binding and subsequent toxicity.

### Combinations of markers could begin to address target expression heterogeneity in HCC

Given the heterogeneity of HCC, identifying novel combinations of the eight candidate markers presents a strategy to identify and treat a greater proportion of HCC tumors. We hypothesized that a select number of gene pairs expressed in high percentage of malignant cells could serve as a more effective, general HCC signature. We observed a marked increase in the percentage of malignant cells that express either gene from all possible combinations of the malignant, single cell differentially expressed gene pairs (**Fig. 3A** red line). Then, we determined that the majority of differentially expressed genes were expressed in ∼60% of malignant cells (**Fig. 3A**, blue line). AMBP was expressed in the highest percentage of malignant cells at 84.4%. When gene pairs were considered, we saw that ∼75% of malignant cells were targeted for most gene pairs.

**Fig. 3.**
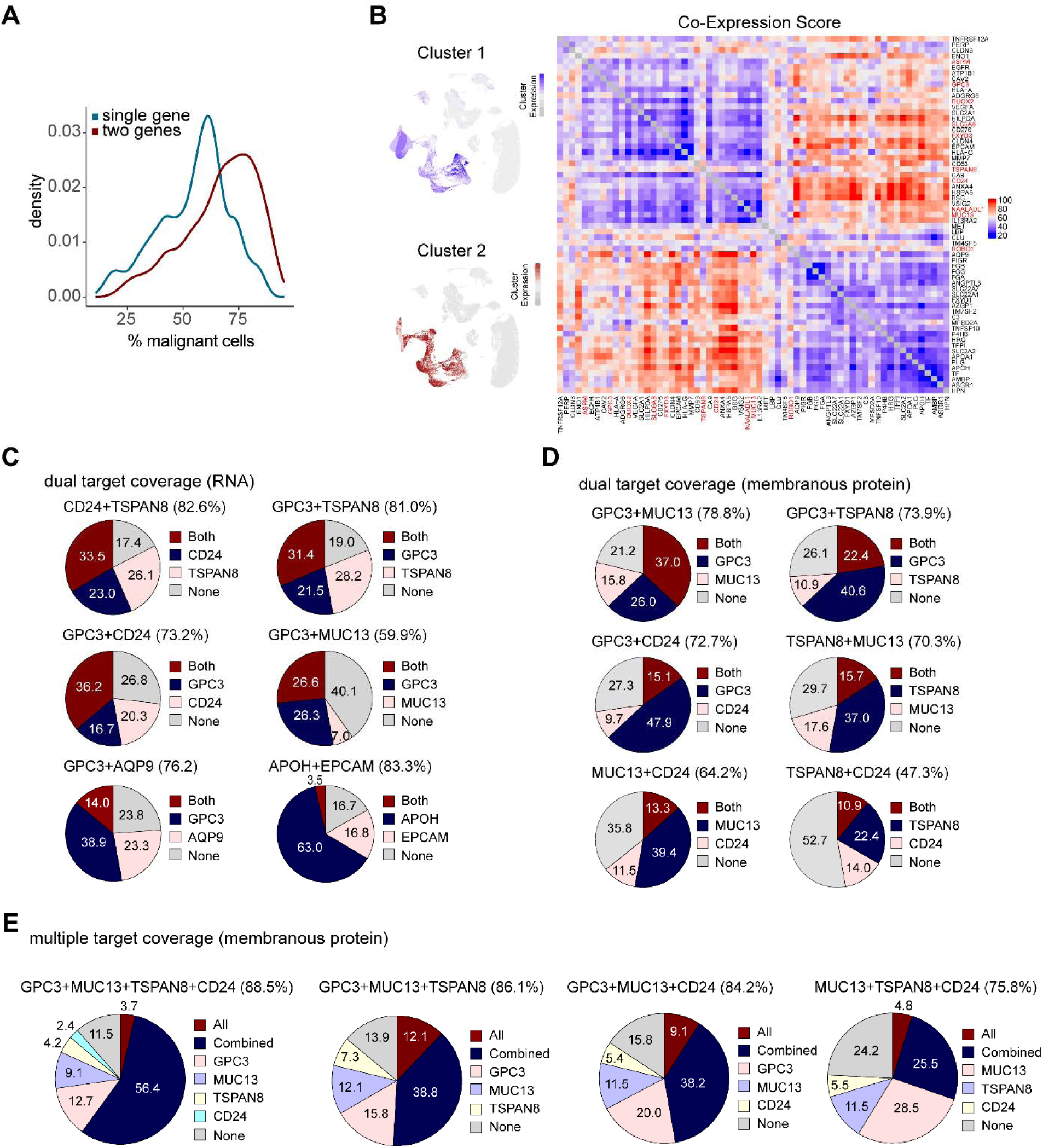
Distribution and coverage of dual and multiple target combinations in HCC based on membrane expression of GPC3, MUC13, TSPAN8, and CD24. (**A**) Distribution of the percentage of malignant cells expressing differentially expressed malignant genes for a single gene (red) or a pair of genes (blue). (**B**) Heatmap of paired Co-Expression scores between differentially expressed malignant genes. Genes are separated into 2 clusters identified by a weighted gene co-expression network analysis (WGCNA). The average gene expression for genes in cluster 1 (blue) and cluster2 (maroon) are projected onto a UMAP of all cells (left). The 10 differentially expressed genes identified in both the single cell and RNA-seq datasets are highlighted in red text. (**C**) Distribution of dual target combinations among select differentially expressed malignant genes based on percentage of malignant cells expressing the gene. The sections of the pie chart indicate the percentage of cells expressing both genes (Maroon), individual genes (blue and pink) or neither gene (grey). (**D**) Distribution of dual target combinations among GPC3, MUC13, TSPAN8, and CD24 in HCC patients based on membrane expression. The pie chart displays the percentage coverage across 165 HCC patients, categorized into both targets, single target, and none. (**E**) Distribution of multiple target combinations among GPC3, MUC13, TSPAN8, and CD24 in HCC patients based on membrane expression. The pie chart illustrates the percentage coverage in the same cohort, categorized into all targets, combinations of two or more targets, single target, and none.

A weighted gene co-expression network analysis (WGCNA) of the malignant, single cell differentially expressed genes identified two clusters of highly correlated gene expression (**Fig. 3B**). The average gene expression in each cluster shows the specificity for malignant cells. A Co-expression score was calculated (see methods) to identify which gene pairs provide the greatest improvement in targeting malignant cells as a percentage compared to an individual gene. Gene pairs within the same cluster have the lowest score and are expressed in similar cells, while gene pairs between clusters have the highest score and show the greatest increase the number of malignant cells over the individual gene expression (**Fig. 3B**). Individual genes with a high score do not necessarily target a high percentage of malignant cells on their own, but when the expression of gene pairs is considered in aggregate, the number of targeted malignant cells greatly increases (**Fig. 3C**).

All shared differential genes between single cell and TCGA data, except ROBO1, are part of the Cluster 1. Individually these genes target between ∼10-50% of malignant cells (**Fig. S2B**). Cluster 1 genes can be used in conjunction with Cluster 2 genes to increase their percentage of targeted malignant cells. This gene pair score can help identify potential targets that may have been previously ignored as they may only target a small percentage of malignant cells individually.

In addition to evaluating combining HCC-selective markers at the transcriptome level, we also assessed at the protein level using IHC. For dual marker combinations based on membrane expression, the combination of GPC3 and MUC13 showed the highest coverage rate of 78.8% (129/165), followed by the combinations of GPC3 and TSPAN8 at 73.9% (122/165), GPC3 and CD24 at 72.7% (120/165), GPC3 and TSPAN8 at 70.3% (116/165), MUC13 and CD24 at 64.2% (106/165), and TSPAN8 and CD24 at 47.3% (78/165) (**Fig. 3D**). Among the multiple marker combinations, the combination of GPC3+MUC13+TSPAN8+CD24 achieved the highest patient coverage rate of 88.5% (146/165). Three-marker combinations demonstrated the following coverage rates: GPC3+MUC13+TSPAN8 at 86.1% (142/165), GPC3+MUC13+CD24 at 84.2% (139/165), and MUC13+TSPAN8+CD24 at 75.8% (125/165) (**Fig. 3E**). Notably, theranostic marker expression in HCC patients exhibits marked heterogeneity (**Fig. S6**).

In aggregate, these findings underscore the potential utility of using multiple marker combinations to address the challenge of heterogeneity of target expression in the development of HCC-selective diagnostic and therapeutic agents.

### Potential theranostic markers are present in various liver cancer cell lines

To rigorously validate whether the HCC-selective molecules were valuable as theranostics targets, we first needed to assess their expression in commercially available cell lines. We performed quantitative real-time PCR (qRT-PCR) to assess mRNA expression levels of eight theranostic markers in five liver cancer cell lines: HepG2, Hep3B, SNU182, SNU449, and Huh7 (**Fig. 4A**). Among the markers analyzed, *CD24* exhibited the highest expression in Huh7 cells, showing a 1.9-fold (*p*<0.05) higher expression compared to Hep3B. Similarly, *MUC13* and *TSPAN8* were predominantly upregulated in Huh7, displaying 52.3-fold (*p*<0.01) and 17.9-fold (*p*<0.01) higher expression levels, respectively, relative to Hep3B. *GPC3* was significantly upregulated in HepG2 cells (4.7-fold higher than Hep3B, *p*<0.0001), whereas SNU182 and SNU449 exhibited minimal or undetectable levels. *ROBO1* expression was comparable between HepG2 and Hep3B, whereas *EGFR* levels were similarly increased in SNU449 and Huh7, with 1.6-fold (*p*<0.05) and 1.9-fold (*p*<0.01) elevations, respectively, compared to Hep3B. *MET* expression varied across cell lines, with SNU449 exhibiting the highest expression (2.2-fold increase, *p*<0.01), while HepG2 and Huh7 displayed comparable upregulation of *BSG*, with 2.1-fold (*p*<0.01) and 2.3-fold (*p*<0.01) increases, respectively, over Hep3B.

**Fig. 4.**
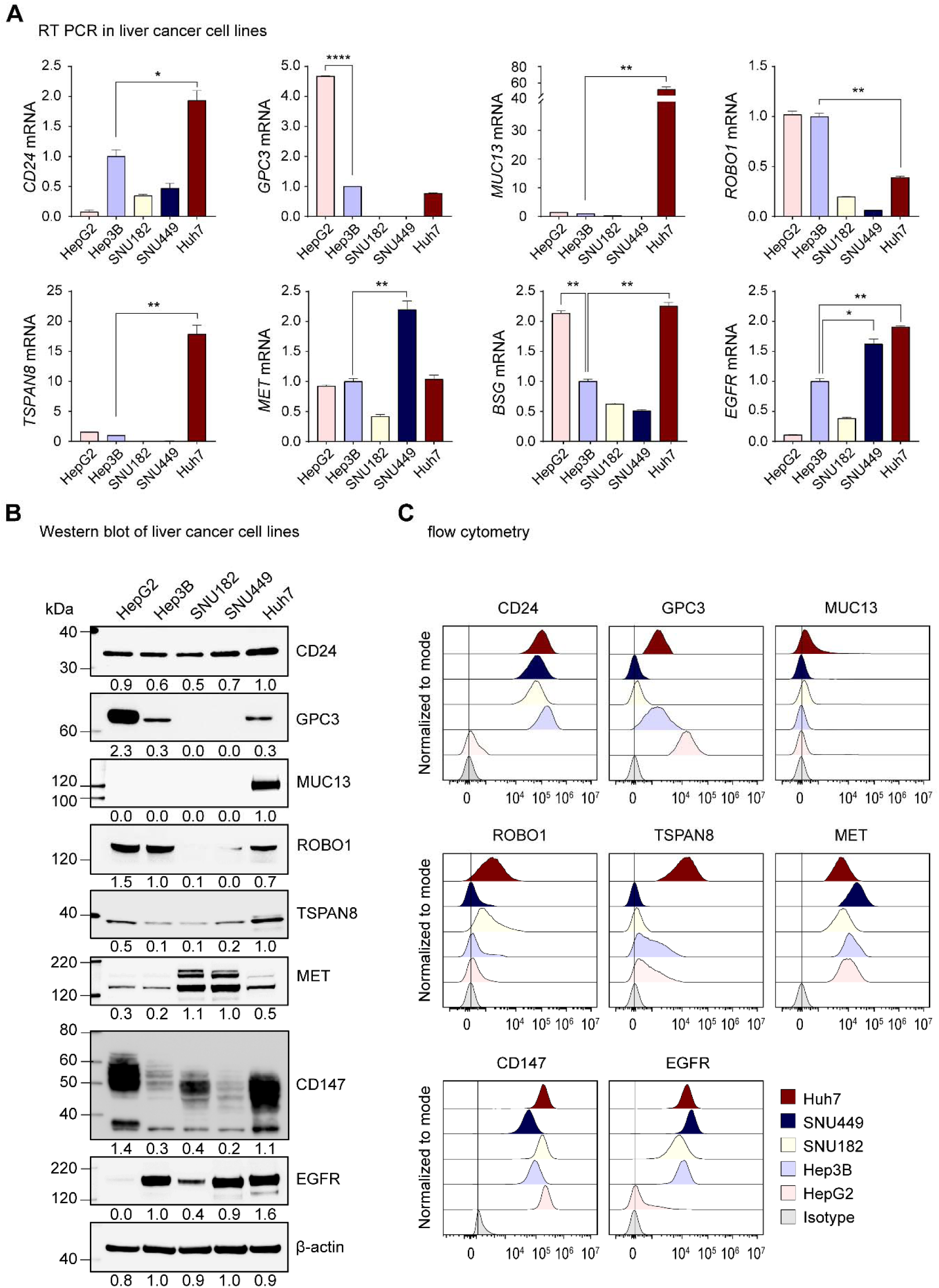
Validation of theranostic targets in liver cancer cell lines. (**A**) Quantitative real-time RT-PCR analysis of the gene expression levels of *CD24*, *GPC3*, *MUC13*, *ROBO1*, *TSPAN8*, *MET*, *BSG* (CD147), and *EGFR* in liver cancer cell lines. *ACTB* served as an internal control, and relative expression levels were normalized to Hep3B. Statistical significance was assessed by two-tailed t-test. *, *p*<0.05; ***, *p*<0.01; ****, *p*<0.0001. (**B**) Western blot analysis of the identified theranostic targets in liver cancer cell lines, with β-actin as the internal control. The numbers beneath the blot images represent the expression levels, shown as fold-change relative to the control. (**C**) Flow cytometry analysis of membranous expression of eight potential theranostic targets across five liver cancer cell lines, with stacked histograms. Solid lines represent cells labeled with the isotype control.

To confirm these findings at the protein level, we performed western blot analysis (**Fig. 4B**), which corroborated the qRT-PCR results. Specifically, GPC3, MUC13, and TSPAN8 exhibited strong expression in HepG2 (2.3-fold), and Huh7 (1.0-fold), respectively, compared to Hep3B. The expression patterns of CD24, MET, and CD147 varied among the liver cancer cell lines. ROBO1 was primarily detected in HepG2, Hep3B, and Huh7, while its expression was negligible in SNU182 and SNU449. EGFR was widely expressed in four HCC cell lines (Hep3B, SNU182, SNU449, and Huh7).

Given the importance of membrane-bound markers for radiotheranostics applications, we further assessed their surface expression. Flow cytometry assays showed high expression of CD24 and EGFR in four HCC cell lines, whereas expression in HepG2 was minimal (**Fig. 4C**). The surface expression patterns of GPC3 aligned with qRT-PCR and Western blot data, showing high levels in HepG2, Hep3B, and Huh7. Additionally, MET and CD147 were consistently expressed on the surface of all five liver cancer cell lines. TSPAN8 was prominently detected on the membrane of Huh7 cells, consistent with its transcriptional expression. Meanwhile, MUC13 demonstrated low surface expression across all tested liver cancer cell lines. These findings show that our HCC-selective markers exhibit distinct expression profiles across liver cancer cell lines, confirming the heterogeneity in liver cancer oncogenesis and the need for rigorous characterization to develop targeted therapeutic interventions.

### Target-selective immunoPET agents successfully localizes to liver cancer xenograft tumors

To assess the potential of the identified markers in this study as theranostic targets for liver cancer, we administered the ^89^Zr- or ^18^F-labeled antibody conjugates and performed immunoPET imaging in several murine models of liver cancer using various cell lines. The imaging results demonstrated excellent tumor-to-background with significant accumulation of the radiolabeled probe at the tumor sites (**Fig. 5A**). Generally, off-target signal was minimal, confirming that utility of diagnostic imaging of the evaluable targets.

**Fig. 5.**
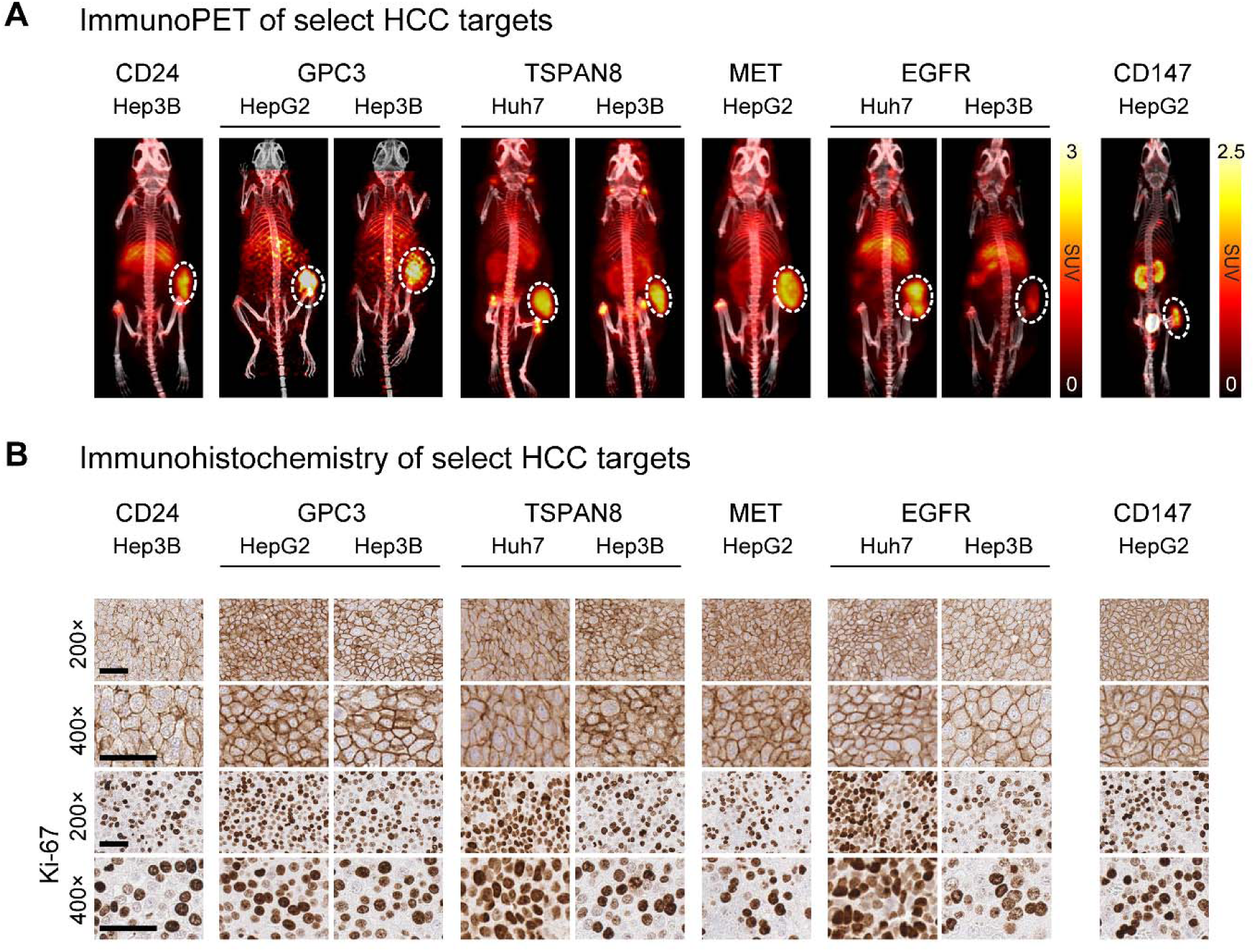
Confirmation of theranostic marker accumulation and expression using ImmunoPET and immunohistochemistry. (**A**) Validation of targeted theranostic marker detection via immunoPET in xenograft models. Representative MIP-PET/CT images of mice bearing liver cancer cell line xenografts, injected with ^89^Zr-DFO-labeled antibodies (anti-CD24, anti-GPC3, anti-TSPAN8, anti-MET, and anti-EGFR) or [¹ F]FPy-DMN1 (for CD147 targeting), and imaged at target-specific time points (2, 72, or 144 hours) post-injection. Tumors are highlighted by white dotted circles. PET images are presented with radioactivity levels calibrated in standardized uptake values (SUV). (**B**) Representative immunohistochemical images of xenograft tissue samples showing the expression of CD24, GPC3, TSPAN8, MET, EGFR, CD147, and Ki-67 following immunoPET. Images are provided at both low (top panels) and high (bottom panels) magnifications. Scale bar = 50 *µ*m.

To assess expression in liver cancer xenografts, we also performed immunohistochemistry analysis (**Fig. 5B)**. Staining results confirmed high membrane expression of the evaluated markers within tumor tissues, demonstrating a positive correlation between both SUV_mean_ (*Pearson’s* ρ=0.707) and SUV_max_ (*Pearson’s* ρ=0.723) and target expression levels by quantified by IHC (**Figs. S7**). Additionally, Ki-67 expression showed similar levels across xenograft tissues derived from liver cancer cells. These promising immunoPET results highlight the potential of these markers to serve as both diagnostic biomarkers and therapeutic targets, paving the way for personalized treatment approaches.

## DISCUSSION

Here, we identify several HCC-selective molecules and demonstrate that they could be used to inform the development of novel diagnostic (e.g. PET) or possibly therapeutic agents (e.g. biomolecule-drug conjugates, radioconjugates, chimeric antigen receptor cellular therapy) using publicly available bulk and single-cell RNA datasets of patients with HCC, validating with patient-derived tissue microarrays of HCC and normal tissues, as well as commercially available liver cancer cell lines. Our approach builds and expands upon prior efforts to identify theranostics targets (*34*) with experimental validation studies. These efforts could allow both the repurposing of existing targeted therapies that are approved for other indications for use in patients with HCC as well as the development of new ones.

Some of the targets we identified have been reported in the literature, however, at the time of this writing, only GPC3- and CD147-targeted agents have been evaluated in HCC clinical trials (*18, 21, 35, 36*). Importantly, the GPC3-targeted antibody, codrituzumab, has previously been studied in a phase 2 clinical trial of patients with advanced HCC randomized to either codrituzumab or placebo (*18*). This trial was negative for its primary endpoint of progression-free survival and secondary endpoint of overall survival. These data suggest that antibody-dependent cellular cytotoxicity, the putative mechanism of action for unconjugated antibodies, is insufficient to exert a therapeutic benefit despite the codrituzumab having a very high affinity for GPC3 and demonstrating good HCC localization in humans (*22*). For such targets, it is not unreasonable to consider coupling a cytotoxic cargo to the targeting biomolecule (e.g., antibody, peptide) and enhancing cytotoxicity for therapy. In the radiopharmaceutical realm, the cargo could be either a radionuclide with diagnostic or therapeutic emissions (*24–27, 37, 38*). To date, there are two GPC3-specific radiopharmaceutical therapy agents, one based on a cyclic peptide (NCT06726161) and another based on a full-length antibody (NCT06764316) are being tested in clinical trials using the therapeutic alpha particle emitter actinium-225.

A CD147-specific F(ab’)_2_, metuximab, labeled with ^131^I is approved in China for the treatment of HCC. In a phase 2 randomized controlled clinical trial, this treatment was shown to improve 5-year relapse-free survival compared to placebo in patients with HCC following surgical resection (*21*). This work underscores the potential of CD147-targeted radiopharmaceutical agents in the treatment of HCC, specifically, and perhaps, the modality’s efficacy more generally. Importantly, this agent is administered transarterially, suggesting that this approach may allow for therapy despite the significant non-tumor tissue expression of CD147. In addition to GPC3 and CD147, we identified several other putative HCC-selective theranostic targets including CD24, TSPAN8, MET, MUC13, and EGFR. While some had been previously described, many were understudied in HCC, especially with respect to radiotheranostics applications.

CD24 is a glycoprotein frequently overexpressed in cancers such as breast, lung, hepatocellular, and ovarian cancers, where it promotes tumor progression, metastasis, and poor prognosis. It drives oncogenesis by regulating key signaling pathways, including Src/STAT3, WNT/β-catenin, and EGFR, and contributes to drug resistance, such as sorafenib resistance in HCC (*39*). CD24 also facilitates immune evasion by interacting with the Siglec-10 axis, making it a critical player in cancer development and progression (*40*). While CD24 has been pursued as therapeutic target for ADCs (*41–43*) and CAR cell therapies (*44, 45*), we are the first to evaluate its potential as a radiotheranostic and reported a more detailed characterization in a separate manuscript (*46*). Importantly, because we observed significant uptake in non-tumor tissues, we advise caution when pursuing CD24 as a therapeutic target for HCC or other malignancies if administered intravenously.

The tetraspanins (TSPAN) are glycoproteins involved in key cellular processes such as migration and growth. Tetraspanin-8 (TSPAN8) is overexpressed in several cancers, including ovarian (*47*), HCC, gastric and colorectal cancer (CRC) (*48*) and is associated with poor patient prognosis. Maisonial-Besset and colleagues successfully demonstrated that their full-length antibody-based radiotheranostics agents ([^111^In]In-Ts29 and [^177^Lu]Lu-Ts29) could specifically localize to and treat preclinical models of CRC (*49*). Here, we confirm that TSPAN8 may also serve as a valuable radiotheranostics target in HCC. A separate report detailing the characterization of the immunoPET agent, briefly included here is currently under revision.

MET, also known as the hepatocyte growth factor receptor (HGFR), is a receptor tyrosine kinase that plays a crucial role in various cellular processes (*50*). Dysregulation of this pathway, including overexpression or activating mutations of MET, is commonly associated with tumorigenesis and metastasis in various cancers, including HCC (*51, 52*). Several imaging agents targeting MET are being developed or studied for MET aberrant cancers utilizing MET overexpression to enhance tumor detection and characterization, however, most studies have focused on non-HCC histologies (*53–55*). Our prior work successfully testing the MET-specific antibody onartuzumab as a targeting vector in models of pancreatic adenocarcinoma as a radiotheranostic agent in that disease, prompted us to assess its utility in HCC after identifying MET as a potential target for the disease. The results presented here suggest that this target could be useful as an imaging biomarker and help to address the gap and improve diagnostic precision.

MUC13, another HCC-selective marker we identified, is a high-molecular-weight transmembrane glycoprotein that is frequently overexpressed in various cancers, including HCC (*56*). Its elevated expression in HCC has been linked to tumor progression and an increased hepatitis B virus copy number (*57*). Furthermore, MUC13 contributes to the activation of the Wnt/β-catenin signaling pathway by facilitating the nuclear translocation of β-catenin, which upregulates downstream effectors like c-Myc and cyclin D (*58*). A recent abstract reported on successful visualization of MUC13 by PET/CT using a ^89^Zr-labeled full length antibody in a xenograft mouse model of CRC (*59*). In our study, we observed that MUC13 expression was higher in HCC compared to normal liver tissue, with minimal cross-reactivity in other normal organs, save for colon and small intestine. These results highlight the potential of MUC13-targeted immunoPET for radiotheranostics applications in HCC. Unfortunately, at the time of this writing, we were unable to secure a good biomolecule specific to MUC13 to perform immunoPET studies.

EGFR is a well-established receptor tyrosine kinase that has been a target for several malignancies, and HCC is no exception. In fact, a phase 3 clinical trial was performed evaluating the then standard-of-care sorafenib, a pan-tyrosine kinase inhibitor, alone versus sorafenib in combination with erlotinib, a small molecule tyrosine kinase (TKI) inhibitor of EGFR, was reported in 2015 (*60*). The results showed that erlotinib did not improve the primary endpoint of survival. Still, given the overexpression of EGFR in HCC, Takao et al. leveraged this feature to successfully deliver EGFR-targeted photodynamic immunotherapy in preclinical models of HCC (*61*). Future work evaluating EGFR as a radiopharmaceutical therapy in preclinical models of HCC is ongoing.

Curiously, while ROBO1, the human homolog for the Drosophila roundabout gene, has been evaluated for radiotheranostic potential and the RNA analyses suggested it could be promising, we concluded it to be less compelling because of its relatively low expression in tumor TMAs and commercially available cell lines (*17*). It is possible that ROBO1 is expressed in a high proportion of HCC albeit at a low level; however, our workflow would not have identified it as a viable target based on our criteria.

Beyond the HCC-expressing targets, we also interrogated the possibility of tumor microenvironment targets, including FOLH1 (a.k.a. PSMA), which is expressed in the neovasculature of some tumors including HCC as well as FAP, which is expressed on fibroblasts, for which there are available radiotheranostics agents that are either approved or being tested clinically. Because tumor-selective imaging would be beneficial in diagnosis, surveillance, and treatment responses monitoring for patients with HCC, both targets are actively being evaluating in clinical studies in patients with HCC (*13, 62–65*).

We should emphasize that the primary purpose of our efforts was to identify and validate the putative HCC targets for diagnostic and therapeutic development, not to present the immunoPET agents generated as candidates for further development themselves. For example, having demonstrated that some of these markers have high expression in normal tissue, one could either decline pursuit of that target or attempt to balance tumor-selectivity and on-target, off tumor potential for toxicity. Thus, the framework presented here can help distinguish targets to be further explored from those that should not.

In conclusion, we have identified and validated several HCC-selective targets that could be leveraged toward novel targeted diagnostic and/or therapeutic agents, ranging from radiotheranostics through antibody-drug conjugates to CAR cell therapy, or, frankly, any approach targeting molecules that are overrepresented on the plasma membrane of tumor cells. One major potential benefit in identifying some of these less common targets is that we could mix-and-match 2-3 targets for treatment to maximize the number of HCC we could treat and begin to address the challenges of tumor heterogeneity especially when engineering agents that are specific to multiple targets. While our efforts are promising for HCC, the workflow could be generalized to other cancer histologies.

## MATERIALS AND METHODS

### Study design

This study was designed to systematically discover, validate, and preclinically qualify tumor-selective, cell-surface targets suitable for radiotheranostic development in HCC. We first performed an integrated multi-omic discovery analysis using TCGA bulk RNA-sequencing and harmonized single-cell RNA-sequencing datasets to identify malignant-cell–enriched, membrane-localized genes with minimal expression across normal hepatic, stromal, and immune compartments. Candidates meeting predefined malignant-cell enrichment and surface-accessibility thresholds were subsequently advanced to protein-level assessment using large-scale human HCC tissue microarrays supplemented with multi-organ normal controls to evaluate tumor selectivity and mitigate potential off-tumor/on-target safety liabilities.

Prioritized markers were then interrogated in multiple human HCC cell lines to confirm transcript abundance and surface protein expression using quantitative RT-PCR, Western blotting, and flow cytometry. Targets demonstrating consistent surface localization and sufficient tumor prevalence progressed to in vivo qualification using xenograft mouse models and zirconium-89 or fluorine-18 radiolabeled antibody- or nanobody-based immunoPET tracers to evaluate tumor uptake, whole-body biodistribution, and translational feasibility. All human and animal studies were approved under institutional ethical oversight, and detailed experimental protocols are provided in the Materials and Methods section of the Supplementary Materials.

### Statistical analysis

Data were analyzed using SPSS Statistics for Windows, version 29 (IBM Corp., Armonk, NY, USA), and Prism 10.1.2 (GraphPad Software Inc., San Diego, CA, USA). Differential expression of tumor-associated malignant cell surface markers was assessed with two-tailed paired t-tests.

Survival was evaluated using Kaplan–Meier curves and compared by log-rank tests. Combinational expression patterns were visualized with pie charts. For RNA-seq data, we quantified the proportions of malignant cells co-expressing gene pairs or expressing either gene alone. Independent predictors of survival were identified using Cox proportional hazards regression. Statistical significance was defined as *p* < 0.05.

## Acknowledgments

This research was fully supported by the Intramural Research Program of the National Cancer Institute, National Institutes of Health (Molecular Imaging Branch, Center for Cancer Research). The contributions of the NIH authors were made as part of their official duties as NIH federal employees, are in compliance with agency policy requirements, and are considered Works of the United States Government. However, the findings and conclusions presented in this paper are those of the authors and do not necessarily reflect the views of the NIH or the U.S. Department of Health and Human Services. We also acknowledge the NIH Cyclotron Facility for the production and supply of the radioisotopes used in this study.

## Author contributions

Conceptualization: F.E.E.

Methodology: P.H., J.Y.C., W.L., D.N., J.S.K., H.M., O.J., A.P.K.

Investigation: P.H., J.Y.C., W.L., S.F., O.J., K.K., J.W.K., S.R.L., M.C., F.E.E.

Visualization: P.H., J.Y.C., W.L., O.J., F.E.E.

Funding acquisition: F.E.E.

Project administration: P.H., J.Y.C., F.E.E.

Supervision: S.M.H., X.W.W., M.H., F.E.E.

Writing – original draft: P.H., J.Y.C., W.L., O.J., F.E.E.

Writing – review & editing: P.H., J.Y.C., W.L., D.N., J.S.K., H.M., S.F., O.J., K.K., J.W.K., S.R.L., M.C., S.M.H., X.W.W., M.H., F.E.E.

## Data and materials availability

All data are available in the main text or the supplementary materials.

