## Supplementary file for "Multimodal Approach for Identification and Validation of Hepatocellular Carcinoma Targets for Radiotheranostics"

**Correspondence to:**

**Freddy E. Escorcia, MD, PhD**

Molecular Imaging Branch, Radiation Oncology Branch, Center for Cancer Research, National Cancer Institute, National Institutes of Health, Bethesda, MD 20892, USA.

ORCID ID: <https://orcid.org/0000-0002-0727-3242>

**SUPPLEMENTARY RESULT**

**Cytoplasmic expression of *in silico*-identified theranostic markers are highly expressed in HCC patient samples and associated with clinical outcomes**

Cytoplasmic expression of these markers, CD24, GPC3, MUC13, ROBO1, MET, and CD147 were significantly overexpressed in HCC compared to non-tumor tissues (all *p*<0.0001), while TSPAN8 displayed a moderate increase (*p*<0.05). EGFR cytoplasmic expression did not show significant differences between HCC and non-tumorous tissues (**Fig. S4A**).

In the analysis of cytoplasmic expression of selected markers in normal tissue microarrays, a comparable pattern of localization was observed. The markers CD24, GPC3, MUC13, ROBO1, TSPAN8, MET, and CD147 demonstrated cytoplasmic distribution patterns that closely mirrored their membranous expression. However, EGFR showed a notably higher cytoplasmic expression in several normal tissue types (**Fig. S4B**). Survival analysis revealed that high cytoplasmic expression of CD24 (*p*=0.003), ROBO1 (*p*=0.024), and CD147 (*p*=0.001) was significantly associated with shorter DFS (**Fig. S4C**). In addition, low MUC13 cytoplasmic expression correlated with poorer DFS, while increased MET cytoplasmic expression was linked to both worse DFS (*p*=0.002) and OS (*p*=0.029) (**Figs. S4C, 4D**). Multivariate Cox analysis identified high ROBO1 cytoplasmic expression as a significant prognostic predictor for DFS (HR=2.367; 95% CI: 1.381–4.055; *p*=0.002). Furthermore, high CD147 cytoplasmic expression was found to be an independent predictor for both DFS (HR=2.036; 95% CI: 1.168–3.547; *p*=0.012) and OS (HR=3.457; 95% CI: 1.097–10.899; *p*=0.034) (**Table S1**).

For combinations based on cytoplasmic expression, dual marker combinations showed HCC coverage rates ranging from 86.1% for TSPAN8 and CD24 to 90.3% for GPC3 and MUC13 (**Fig. S5A**). Additionally, the combination of multiple markers based on cytoplasmic expressions resulted in HCC patient coverage rates ranging from 95.2% to 98.8% (**Fig. S5B**). Similarly, the combination of dual and multiple markers that incorporate both membranous and cytoplasmic expressions showed comparable HCC patient coverage, with the highest coverage rates of 90.3% for dual marker combinations (**Fig. S5C**) and 98.8% for multiple marker combinations (**Fig. S5D**).

**MATERIALS AND METHODS**

To integrate bulk and single cell RNA sequencing, tissue microarray, cell line and in vivo studies, we used the workflow illustrated in **Fig. S8**.

**TCGA Bulk RNA sequencing analysis**

Bulk RNA-seq data and associated clinical information were obtained from The Cancer Genome Atlas (TCGA). HTseq expression data for LIHC were obtained from the TCGA-LIHC dataset (*66*). Data analysis and visualization were performed within the NIH Integrated Data Analysis Portal (NIDAP) using R programs developed on the Foundry platform (Palantir Technologies, Denver, CO). Expression data from TCGA- LIHC generated by HTseq were transformed to CPM counts, and genes with CPM values less than 6 in at least 50 samples were removed. Differentially expressed genes between tumor and Normal samples were identified using the voom algorithm (*67*) from the Limma R package (version 3.40.6) (*68*). Genes with adjusted *p*-value ≤ 0.005 and log_2_ fold change of ≥ 1.0 were considered significantly differentially expressed.

**Single Cell RNA sequencing dataset analysis**

scRNA-seq data processed using Seurat v4 (*69*). Datasets were filtered to remove low quality cells identified by cell expressing less than 600 genes, total UMIs less than 700, mitochondrial expression greater than 20% and cell complexity (log_10_(Genes/UMI)) less than 0.5. The remaining cells were normalized using SCTransform. Batch correction between samples was corrected using the Seurat anchor-based workflow on 3000 variable features and integration using the CCA algorithm. After Integration Principal Component Analysis (PCA) and UMAP dimensionality reduction was performed.

Cell type classification was assigned using SingleR (*70*) with a reference created from the Human Single Cell Atlas (*32*). The malignant cell classification from Lichun Ma et al. were identified using InferCNV and described in their paper (*71*). Cells from GSE149614 that clustered with the previously annotated malignant cells from GSE125449 were also classified as malignant. Malignant cell markers were identified using the MAST (*72*) in Seurat, comparing malignant cells to all other cell populations. Malignant cell modules were identified using high dimensional WGCNA (*73-75*) and heat map was created using correlation of gene expression across all cells. Gene pairs were colored based their co-expression score calculated as shown below:

$$\left( A_{unique}+B_{unique} \right)\frac{{\%Malignant}_{comb}}{{\%Malignant}_{ind}}$$

A higher co-expression score for a gene pair indicates that individual genes target different malignant cell populations and combine to target larger percentage of malignant cells compared to an individual gene.

**Cell culture**

Human hepatocellular carcinoma (HCC) cell lines (Hep3B, SNU182, and SNU449) and a hepatoblastoma cell line (HepG2) were obtained from the American Type Culture Collection (ATCC; Manassas, VA, USA). The Huh7 HCC cell line was obtained from Dr. Mitchell Ho (Bethesda, USA). All cells were tested for mycoplasma using the Mycoplasma Detection Kit (Thermo Fisher Scientific, San Jose, CA, USA) and grown in Dulbecco’s modified Eagle’s medium (DMEM; Thermo Fisher Scientific), Roswell Park Memorial Institute (RPMI) 1640 medium (Thermo Fisher Scientific), or Eagle’s minimal essential medium (EMEM, ATCC) supplemented with 10% fetal bovine serum (Thermo Fisher Scientific) at 37°C under 5% CO_2_ incubator/humidified chamber (PHCbi, Wood Dale, IL, USA).

**Quantitative RT-PCR**

Total RNA (1 µg) was extracted using the RNeasy Micro Kit (Qiagen, Valencia, CA, USA) and reverse transcribed with the QuantiTect reverse transcription kit (Qiagen), according to the manufacturer’s protocol. Real-time PCR reactions were performed with 50 ng of cDNA in a total volume of 20 µL, using TaqMan® Universal PCR Master Mix (Applied Biosystems, Foster City, CA, USA) on an ABI 7500 system (Applied Biosystems). Predesigned and labelled primer/probe sets were used for following genes: BSG (Hs00936295_m1), EGFR (Hs01076090_m1), GPC3 (Hs01018936_m1), TSPAN8 (Hs00610327_m1), MET (Hs01565584_m1), MUC13 (Hs00217230_m1), ROBO1 (Hs00268049_m1), and CD24 (Hs02379687_s1). Relative mRNA expression levels were determined using the comparative cycle threshold (2^−ΔΔCt^) method, with *ACTB* as the endogenous control, and were subsequently normalized to the Hep3B cell line.

**Western blot analysis**

Cells (5 × 10^6^) were harvested and lysed and extracted with Pierce^TM^ RIPA buffer (Thermo Fisher Scientific) supplemented with Halt^TM^ Protease Phosphatase Inhibitor Cocktail (Thermo Fisher Scientific). Protein concentration was determined using the Pierce^TM^ BCA Protein Assay kit (Thermo Fisher Scientific) according to the manufacturer’s instructions. Proteins (20-50 μg) were resolved using 4-12% sodium dodecyl sulphate-polyacrylamide gel electrophoresis (SDS-PAGE) and then transferred onto a nitrocellulose membrane (Thermo Fisher Scientific) using an electric transfer system (Thermo Fisher Scientific). The membranes were incubated overnight at 4 ℃ with primary antibodies: Anti-Basigin/EMMPRIN (#13287), anti-Met/pre-Met (#8198), anti-MUC13 (#44454), and anti-EGF Receptor (#4267), all diluted 1:1000, obtained from Cell Signaling Technology (Danvers, MA, USA); anti-TSPAN8 antibody (ab70007; dilution 1:1000; Abcam, Cambridge, MA, USA); anti-CD24 antibody (SWA11 clone; dilution 1:1000; Cell science, Newburyport, MA, USA); anti-GPC3 antibody (1G12 clone; dilution 1:1000; Cell Marque, Rocklin, CA, USA); and anti-ROBO1 antibody (22D5 clone; dilution 1:1000; Thermo Fisher Scientific). Anti-β-actin (Cell Signaling Technology) was used as the loading control. Subsequently, the membrane was then incubated with horseradish peroxidase-conjugated anti-mouse (#32430) or anti-rabbit (#32460) (Thermo Fisher Scientific) secondary antibodies, and the immunoreactive bands were visualized using Clarity^TM^ Western ECL Substrate (#1705061) or Clarity Max^TM^ ECL Substrate (#1705062) (Bio-Rad, Hercules, CA). The bands were then detected using a ChemiDoc MP imaging system (Bio-Rad).

**Flow cytometry**

Approximately 5 × 10⁵ HCC (Hep3B, SNU182, SNU449, and Huh7) and hepatoblastoma (HepG2) cells were harvested and washed with ice-cold FACS buffer (1% bovine serum albumin in 1x PBS). The cells were incubated with an Fc block (Miltenyi Biotec, cat. no. 13005990, 1:50) on ice for 15 minutes, followed by incubation with monoclonal antibodies (**Table S2**) for 45 minutes at 4°C in FACS buffer. After washing with FACS buffer, corresponding fluorescence-conjugated secondary antibodies (**Table S2**) were added and incubated for 15 minutes at 4°C in the dark. The labeled cells were then washed, and cell-associated fluorescence signals were evaluated using a flow cytometer (Cytoflex, Beckman Coulter). Data were analyzed using FlowJo software v.10.8.1 (FlowJo LLC, Ashland, OR, USA).

**Tissue samples and Immunohistochemistry**

Tissue samples were prospectively collected via surgery from patients who were admitted to the Kangbuk Samsung Hospital or the Kangnam Sacred Heart Hospital between 2010 and 2021. Tumor tissue samples from 165 patients with primary HCC and 165 matched non-adjacent normal epithelium tissues were constructed into Tissue microarrays (TMAs). All the procedures were conducted according to the ethical guidelines of the Declaration of Helsinki, and the study protocol was approved by the Institutional Review Board at Kangbuk Samsung Hospital (IRB No. 2022–10-052-001, Seoul, South Korea) and Kangnam Sacred Heart Hospital (IRB No. HKS2022-12-015, Seoul, South Korea). Additionally, multiple human normal organ TMAs, consisting of 33 organs/99 cores, were purchased from US Biomax, Inc. (Cat. # FDA999L246, Rockville, MD).

Immunohistochemistry (IHC) was performed on formalin-fixed, paraffin-embedded human HCC, human normal organ, or murine xenograft tissues using EnVision+ Dual Link System-HRP (DAKO, Carpinteria, CA). A mouse-on-mouse IHC kit (M.O.M. kit, Vector Laboratories, Newark, CA, USA) was used to detect CD24 expression in murine xenograft tissues according to the manufacturer’s protocol. Detailed immunohistochemistry conditions are outlined in **Table S3**. The antigen-antibody reaction was visualized with DAB+ (3,3-diaminobenzadine; DAKO) for 10 min. TMA sections were lightly counterstained with Mayer’s hematoxylin and examined under light microscopy. Negative control immunoglobulin G (IgG) was used in place of a primary antibody to evaluate nonspecific staining, and appropriate positive control specimens were included within the TMA. The stained TMA slides were scanned using a NanoZoomer 2.0 HT (Hamamatsu Photonics K.K., Hamamatsu City, Japan) at 20x objective magnification. Digital analysis of the images was performed using Visiopharm Integrator System v6.5.0.2303 (VIS; Visiopharm, Hørsholm, Denmark). Briefly, individual digital images of TMA cores were extracted from the whole slide images. The intensity of brown cytoplasmic and membranous staining for potential theranostic markers was quantified using a predefined algorithm and optimized settings, as previously described (*76*). The average intensity of staining from the two tumor cores was calculated as the final value for membranous and cytoplasmic expression.

**Murine xenograft model**

All animal procedures were conducted in accordance with institutional guidelines under a protocol approved by the Institutional Animal Care and Use Committee at the National Institutes of Health. HepG2, Hep3B, and Huh7 cells (5 × 10^6^ cells per mouse) were subcutaneously inoculated into the right flank of female athymic nu/nu mice (7–10 weeks old; Charles River Laboratories, Wilmington, MA). Tumors were allowed to grow to approximately 200 mm³ prior to immunoPET studies.

**Radioconjugate synthesis and quality assurance**

A panel of antibodies targeting human tumor-associated antigens was used for radiolabeling. These included codrituzumab (GC33; anti-GPC3 IgG1, Chugai Pharmaceutical Co., Ltd, Tokyo, Japan), panitumumab (anti-EGFR IgG2κ, Amgen, Inc., Thousand Oaks, CA, USA), α-hTSPAN8 (rat anti-human IgG2B, #MAB4734, Bio-Techne, Minneapolis, MN, USA), onartuzumab (monovalent humanized anti-MET chimeric OA5D5 antibody, Genentech, San Francisco, CA, USA), α-hCD24 (anti-human CD24, supplier undisclosed), and an in-house anti-CD147 nanobody (DMN1). Antibodies were conjugated with a 2.5–3-fold molar excess of *p*-isothiocyanatobenzyl-desferrioxamine (DFO-Bz-NCS; Macrocyclics, Inc.) using an established protocol (*55*). Radiolabeling with zirconium-89 ([^89^Zr]Zr-oxalate; NIH Cyclotron Facility, Bethesda, MD, USA) was carried out by combining 37–130 MBq of [^89^Zr]Zr-oxalate (diluted in 0.5 M HEPES, pH 7.1–7.3) with DFO-antibody conjugates and 10 µL of 2,5-dihydroxybenzoic acid (5 mg mL⁻¹), followed by pH adjustment using 2 M Na₂CO₃. Reactions were incubated at 37 °C for 1 h and challenged with 10 µL of 100 mM EDTA to verify complex stability. Radioconjugates were purified using PD-10 columns (Cytiva, Marlborough, MA, USA) with phosphate-buffered saline (PBS). Radiochemical purity and labelling efficiency were assessed by radio-instant thin-layer chromatography (radio-iTLC) using iTLC-SG strips (Varian, Lake Forest, CA, USA) with 50 mM EDTA in 100 mM ammonium acetate (pH 5.5) as the mobile phase and analyzed on an AR-2000 radio-TLC scanner (Eckert & Ziegler, Wilmington, MA, USA).

For fluorine-18 radiolabeling, the DMN1 nanobody was labelled via a two-step method as described previously (*77*). [^18^F]FPy-TFP was synthesized on a Sep-Pak cartridge at ambient temperature and reacted directly with DMN1 in phosphate buffer (pH 9.2) at 37 °C for 15–20 min. The crude product was purified using a PD-10 column with saline. The final [^18^F]F-DMN1 product showed >99% radiochemical purity with no detectable aggregation, confirmed by analytical size-exclusion HPLC (TSKgel SuperSW3000, Tosoh Bioscience). The mobile phase comprised 0.1 M sodium phosphate, 0.1 M sodium sulfate, 0.05% sodium azide, and 10% isopropanol (pH 6.8), run at 0.35 mL min⁻¹. Radiochemical yields were 12–18% (decay-corrected, n = 5), with specific activities ranging from 100 to 300 Ci mmol⁻¹. Total synthesis time was approximately 45 min.

***In vivo* molecular imaging**

Mice bearing subcutaneous tumors were intravenously administered 3.7–7.4 MBq (5–10 μg) of ^89^Zr-labeled conjugates or [¹⁸F]FPy-DMN1 (for CD147 targeting) via tail vein injection. ImmunoPET imaging was performed at target-specific time points: 72 hours post-injection for GPC3 (HepG2 and Hep3B), EGFR (Huh7 and Hep3B), MET (HepG2), and CD24 (Hep3B ; 144 h for TSPAN8 (Huh7, Hep3B); and 2 h for CD147 (HepG2). GPC3 and EGFR imaging was conducted on a BioPET/CT scanner (BioScan Inc., Washington, DC, USA), whereas TSPAN8, MET, CD24, and CD147 imaging used an MRS*PET/CT 120 scanner (MR Solutions, Guildford, UK). Data acquisition employed VISTA (Sedecal, Madrid, Spain) and Preclinical Scan (v4.2.4.0) software (version 4.2.4.0, MR Solutions). Mice were anesthetized with 2% isoflurane, and static PET scans (10–20 min) were acquired, followed by CT for attenuation correction and co-registration. PET data were reconstructed using 3D OSEM and normalized, decay-corrected, and dead-time–corrected. Images were analyzed using MIM software (version 7.2.7, MIM Software Inc., Beachwood, OH, USA).

References and Notes

References 66–77 are cited only in the Supplementary Materials.

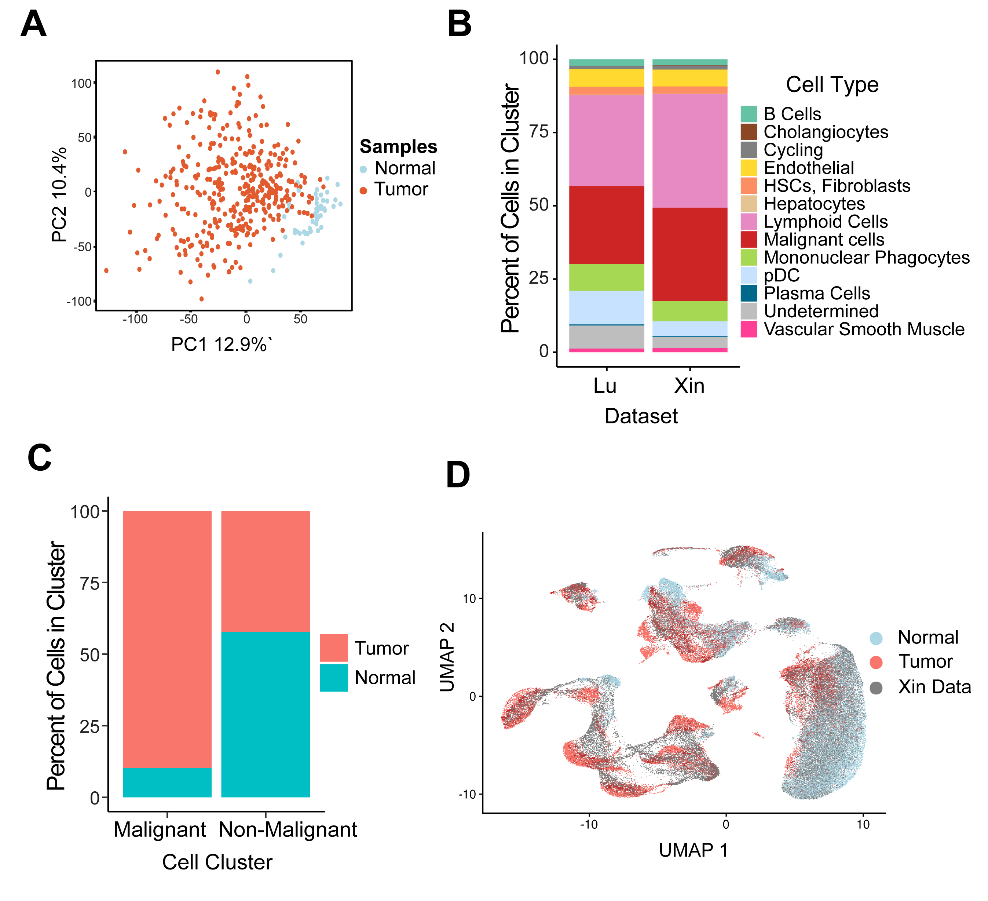

**Fig. S1. Integration of liver tissue Single Cell datasets.** (**A**) PCA plot of bulk RNA seq data from TCGA-LIHC. 371 HCC tumor (red) and 50 normal liver tissue (blue) samples. (**B**) Bar graph for proportion of major cell types in each single cell dataset. (**C**) Bar graph for proportion of malignant and Non-malignant cells in each single cell dataset. (**D**) UMAP projection of all cells from integrated single cell datasets colored by Normal (blue) and Tumor (red) cells.

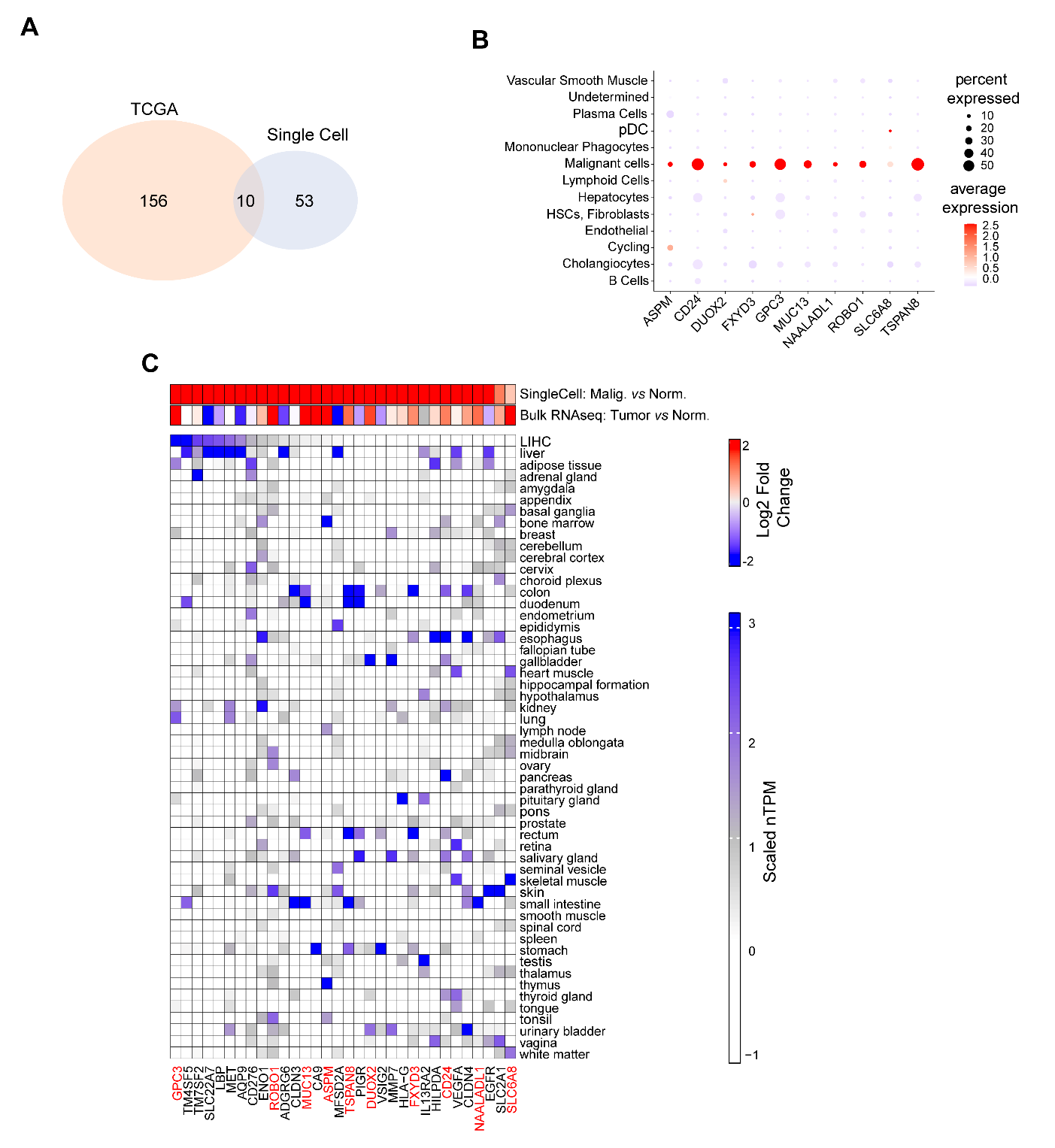

**Fig. S2. Identification and Exploration of HCC specific theranostic targets.** (**A**) Venn Diagram comparing differentially expressed genes identified from the single cell data sets (blue - malignant vs non-malignant) and the TCGH-LIHC bulk RNA-seq data set (red – Tumor vs Normal). (**B**) Dot plot of differentially expressed genes identified in both the single cell and RNA-seq datasets for all major cell populations in the single cell data. The size of the circle indicates the percentage of cells expressing a given gene within the cell type and the color indicates the average expression of the gene (low – white, high – red) (**C**) Heatmap of differentially expressed malignant genes from single cell datasets. Lower heatmap is colored by nTPM values for the consensus dataset and the TCGA cancer dataset (LIHC) from the Human Protein Atlas database (PADB). Genes expression is column normalized across all tissues ranging from low (white) to high (dark blue). The upper heatmap shows the log_2_ Fold Change values from the Single cell dataset comparing Malignant cells to normal cells and the log_2_ Fold Change (blue-decreased to red-increased) from TCGA-LIHC Bulk RNAseq analysis comparing Tumor and normal samples.

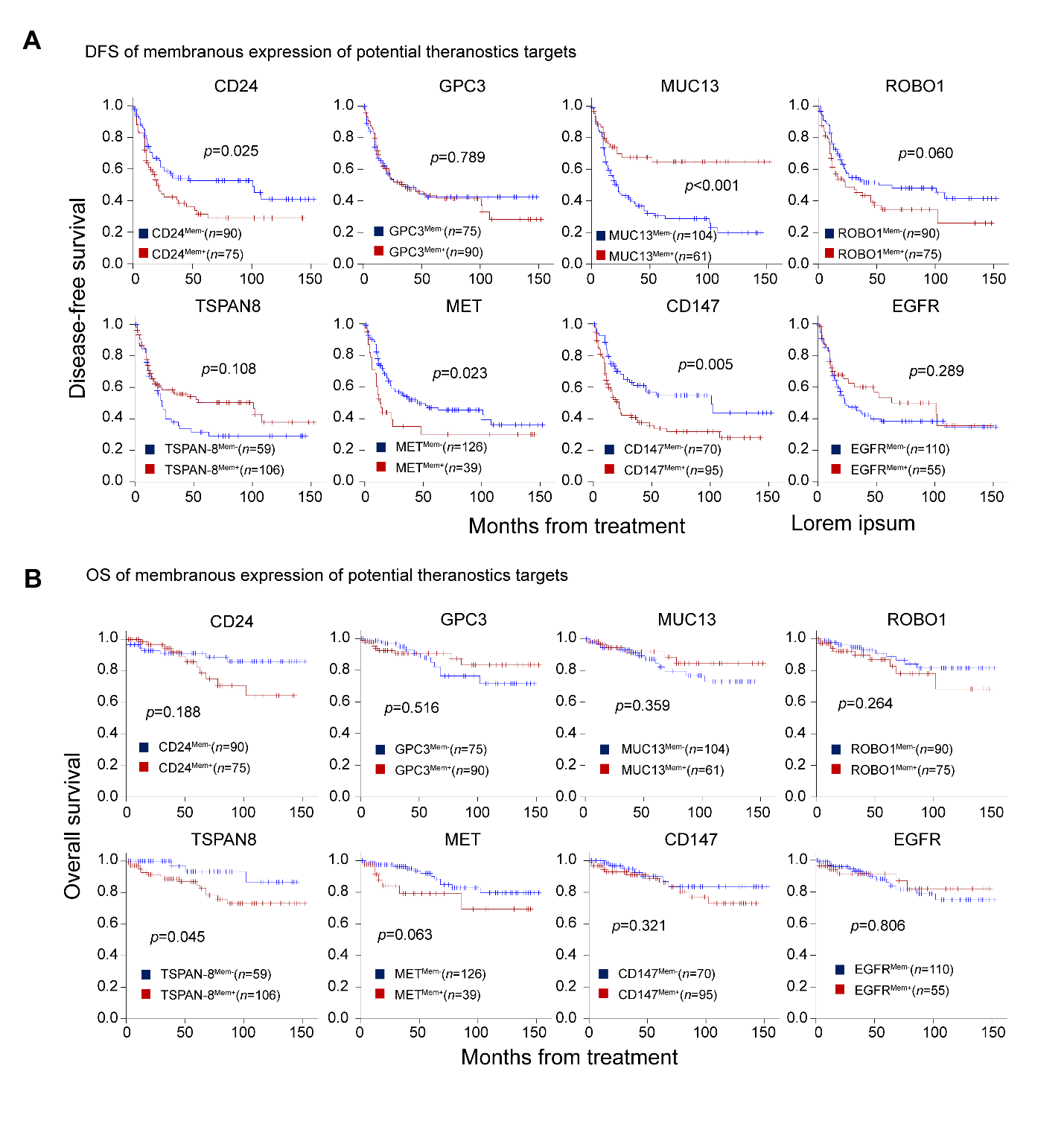

**Fig. S3. Survival analysis of HCC patients based on membranous expression of potential theranostic targets.** Kaplan-Meier survival curves comparing disease-free survival (**A**) and overall survival (**B**) between groups classified by high and low membranous expression levels of theranostic targets.

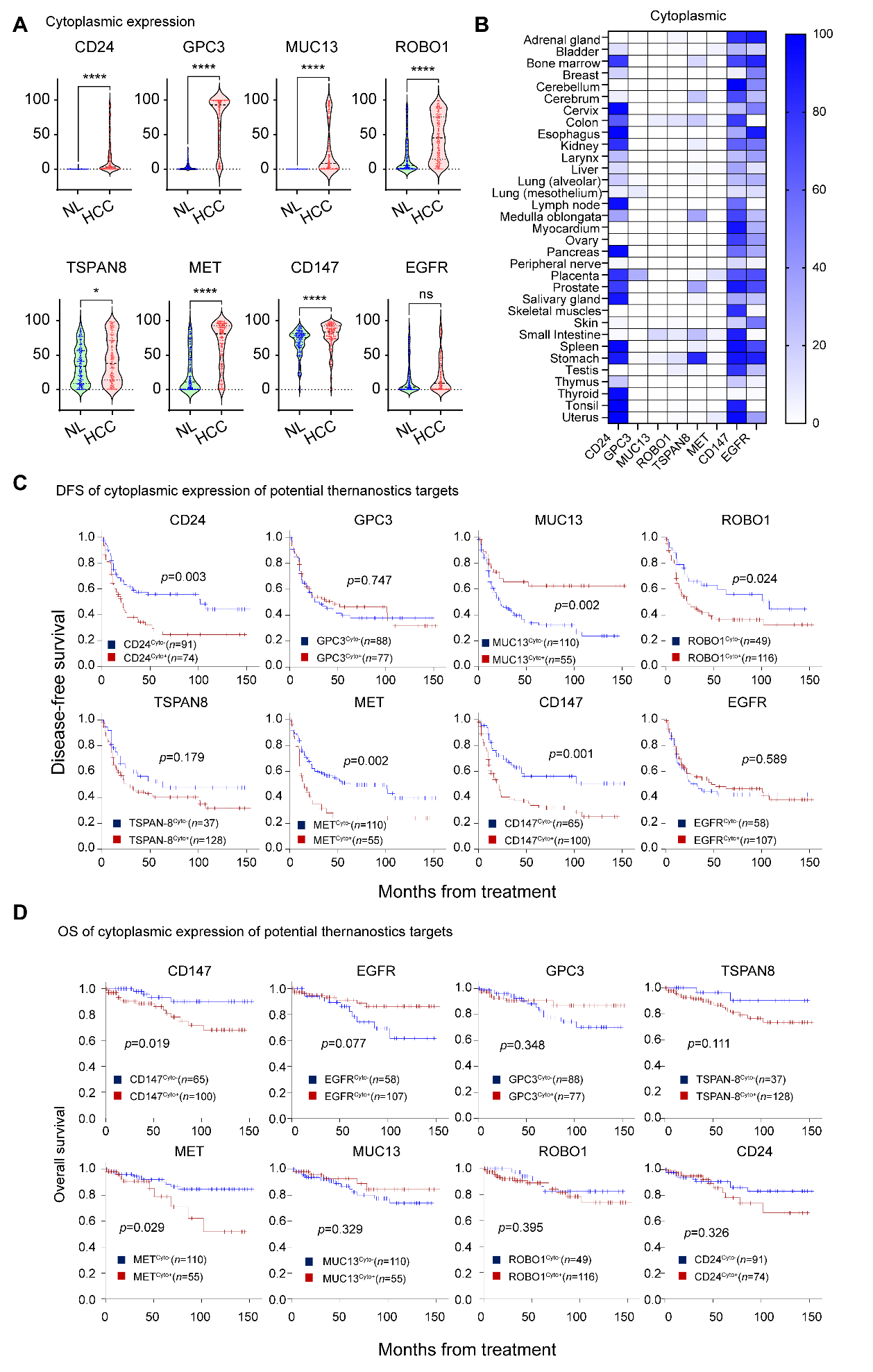

**Fig. S4. Clinical significance of cytoplasmic expression of potential theranostic markers in HCC patients.** (**A**) Violin plots showing the distribution of theranostic markers based on cytoplasmic expression in HCC patient samples and adjacent non-tumor tissues. Statistical analysis was performed using a two-tailed t-test. *, *p*<0.05; ****, *p*<0.0001. (**B**) Heatmap showing protein expression profiles in normal human tissues based on cytoplasmic localization. Kaplan-Meier plots for disease-free survival (**c**) and overall survival (**d**) based on cytoplasmic expression levels of potential theranostic markers in HCC patients.

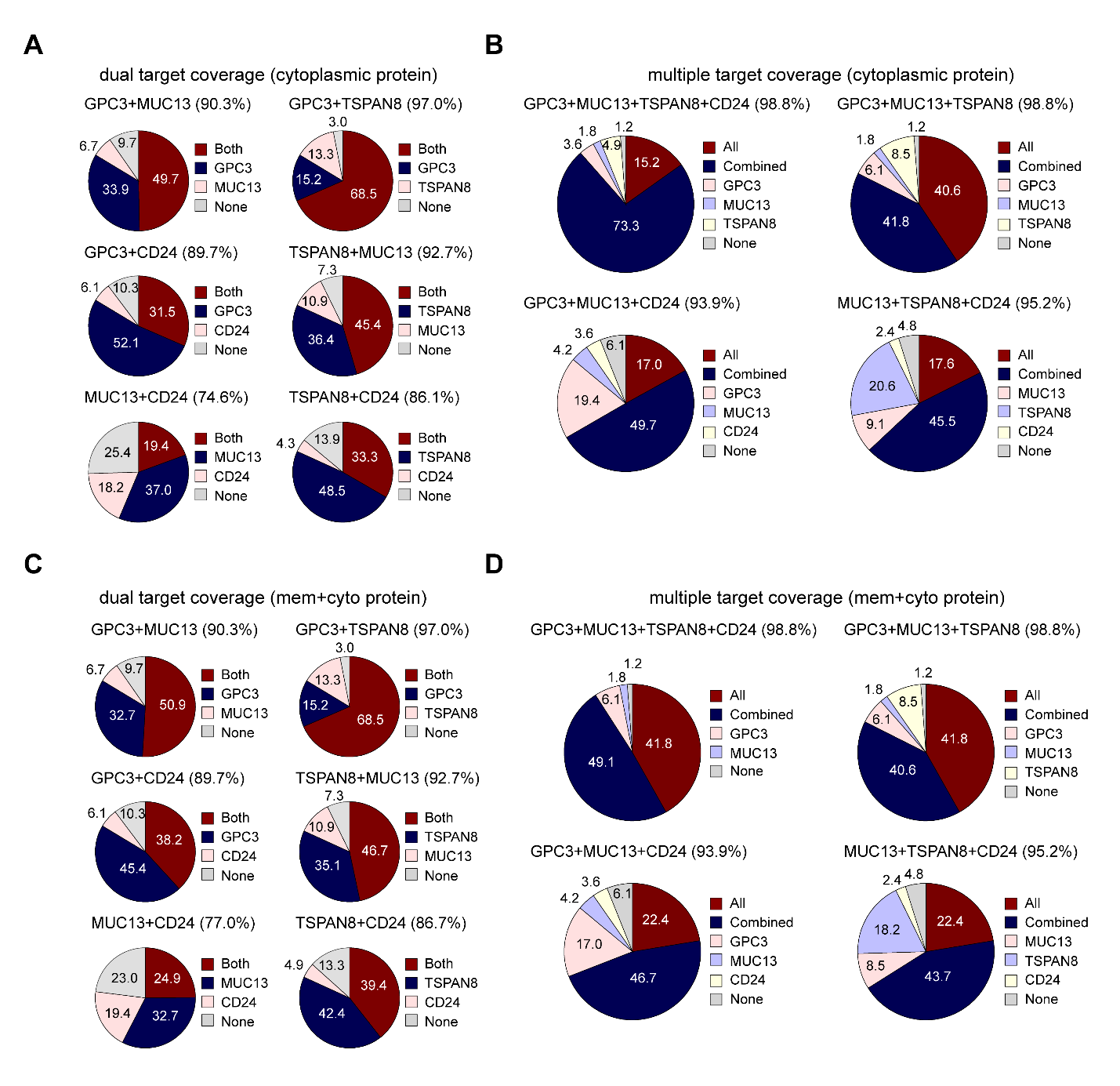

**Fig. S5. Distribution of potential theranostic marker expression in HCC based on cytoplasmic and combined membranous and cytoplasmic localization.** The pie chart (**A**) represents the expression of dual targets in the cytoplasm, while chart (**C**) shows the combined membranous and cytoplasmic expression in 165 HCC patients. The pie charts in (**B**) and (**D**) depict the expression profiles when multiple targets are combined, with coverage percentages indicated on each chart.

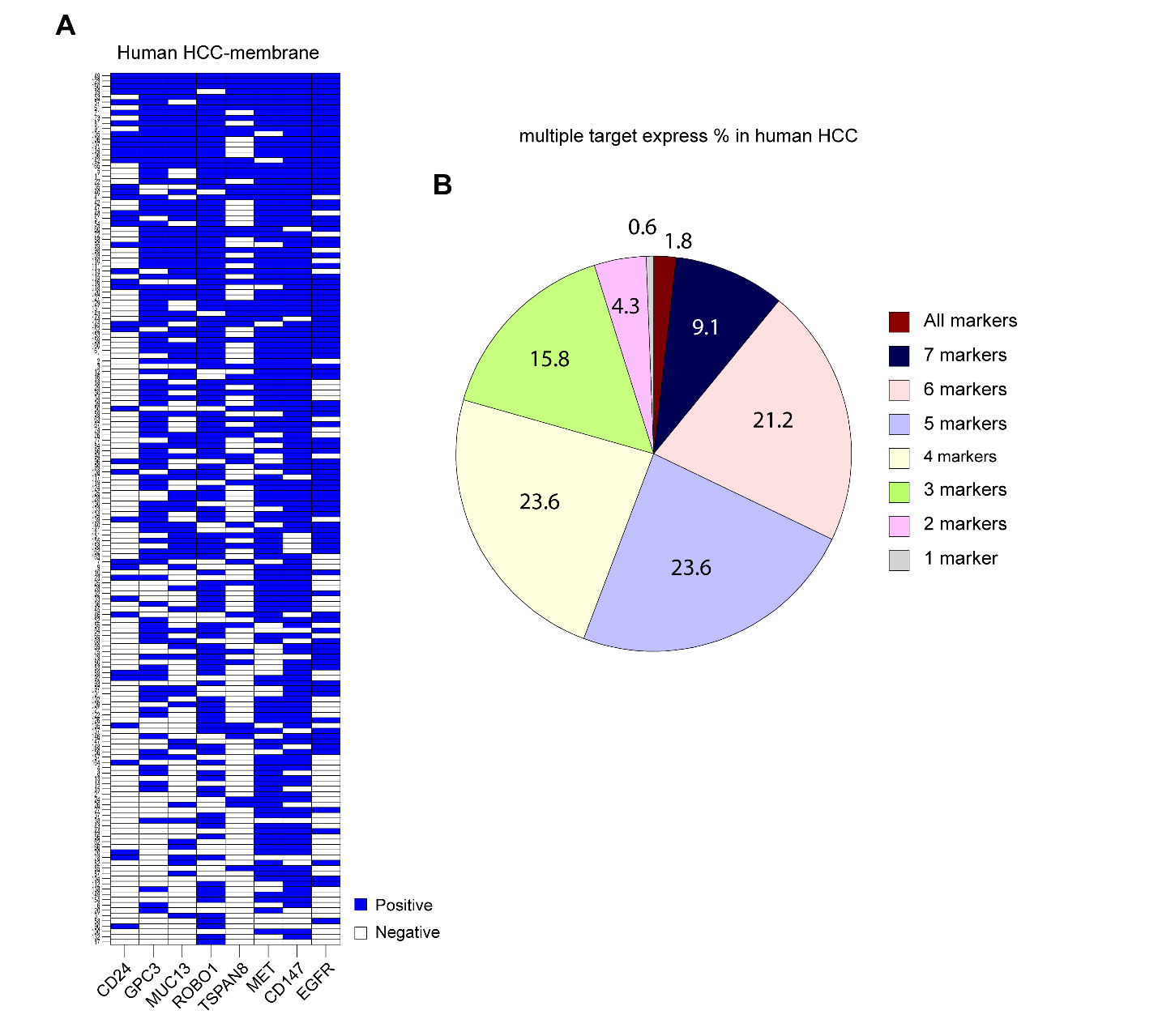

**Fig. S6. Membranous localization and expression patterns of candidate theranostic markers in HCC patient tissues.** (**A**) Map illustrates immunohistochemically positive and negative staining profiles across the HCC tissue cohort. (**B**) Pie chart shows the proportions of distinct combinational theranostic markers expression patterns identified within the patient samples.

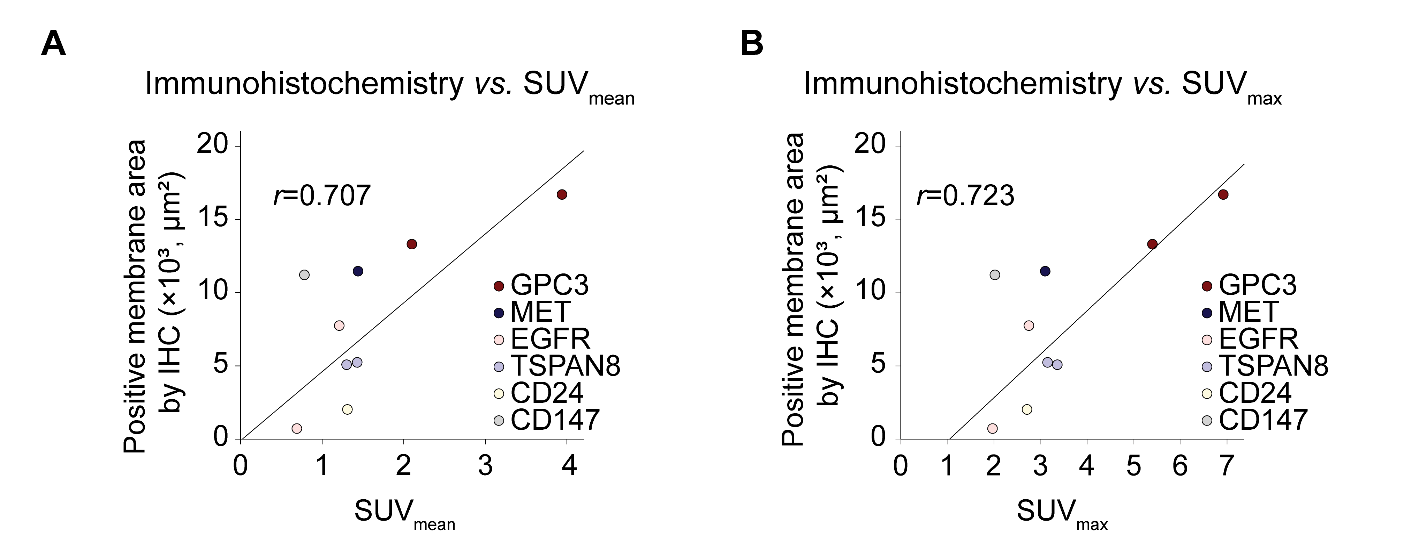

**Fig. S7. Correlation between immunohistochemical membrane staining and PET-derived standardized uptake values (SUVs) in HCC tissues.** Quantitative analysis reveals a strong positive correlation between the extent of membranous immunohistochemical staining and both the mean of SUV (**A**, SUV_mean_, *r*=0.707) and maximum of SUV (**B**, SUV_max_, *r*=0.723).

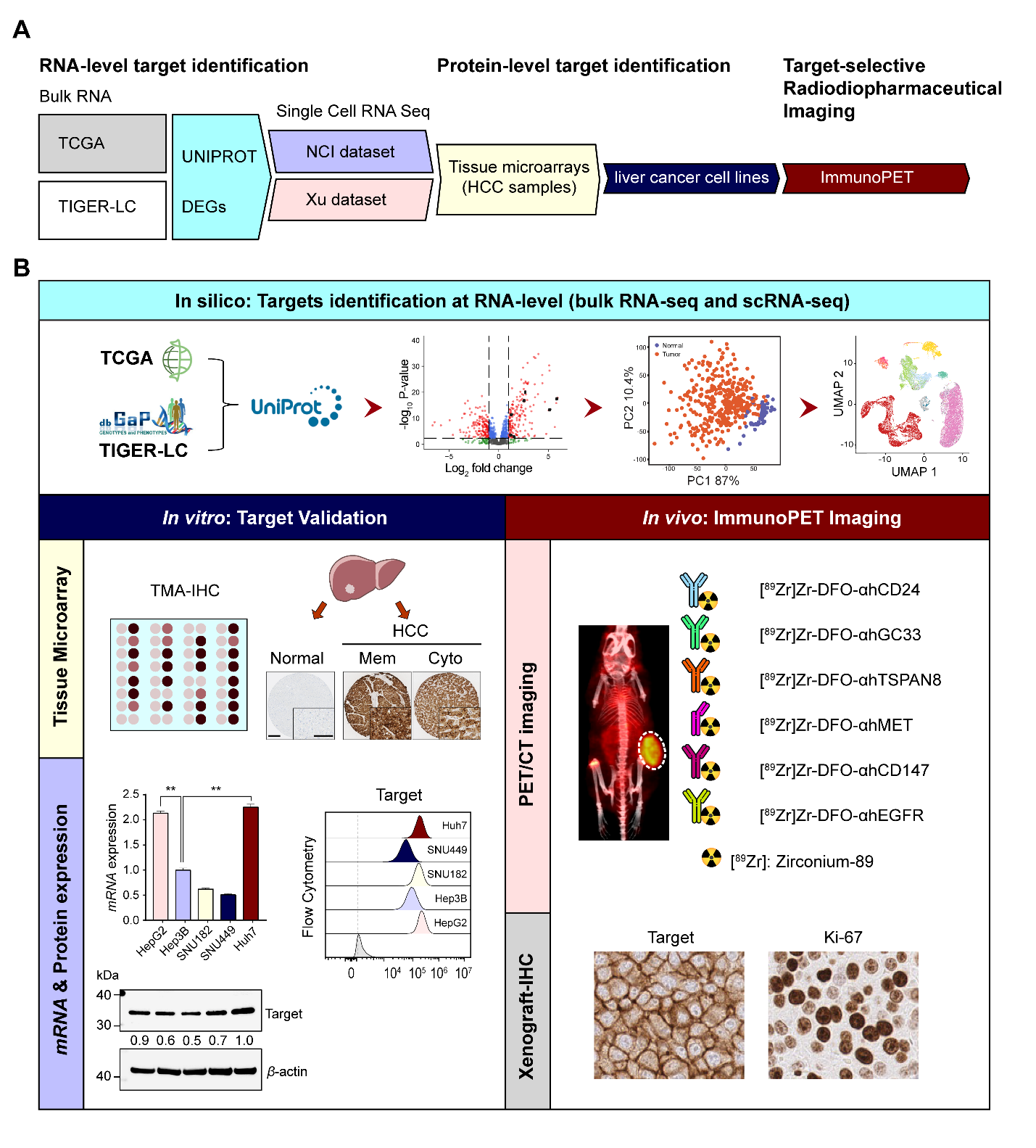

**Fig. S8. Novel radiotheranostic target identification and characterization using an in-silico screening approach and orthogonal *in vitro* and *in vivo* validation assays.** (**A**) Overall workflow of target identification from genome and protein databases. (**B**) Graphical representation of HCC-specific radiopharmaceutical target identification: *in-silico* screen of bulk RNA-sequencing and single cell RNA-sequencing for target generation, *in vitro* target validation at the genomic, protein and *in vivo* level with radio-immunoPET.

**Table S1.** Cox proportional univariate and multivariate analyses of disease-free survival and overall survival in patients with HCC

| **Variables** | **Disease-free survival** | | | | |  | **Overall survival** | | | | |
| --- | --- | --- | --- | --- | --- | --- | --- | --- | --- | --- | --- |
|  | Univariate analysis | |  | Multivariate analysis | |  | Univariate analysis | |  | Multivariate analysis | |
|  | HR [95% CI] | *p* value |  | HR [95% CI] | *p* value |  | HR [95% CI] | *p* value |  | HR [95% CI] | *p* value |
| Age (> 60) | 0.976 [0.635-1.498] | 0.910 |  |  |  |  | 1.461 [0.603-3.540] | 0.402 |  |  |  |
| Sex (male) | 1.677 [0.944-2.979] | 0.078 |  |  |  |  | 0.954 [0.345-2.640] | 0.928 |  |  |  |
| BMI (≥ 25) | 0.889 [0.573-1.379] | 0.598 |  |  |  |  | 0.563 [0.204-1.549] | 0.266 |  |  |  |
| HBsAg^+^ | 0.878 [0.557-1.385] | 0.576 |  |  |  |  | 0.502 [0.203-1.241] | 0.136 |  |  |  |
| pT | 1.575 [1.194-2.078] | 0.001^*^ |  | 1.834 [1.361-2.472] | <0.001^*^ |  | 2.217 [1.275-3.855] | 0.005^*^ |  | 2.700 [1.503-4.848] | 0.001^*^ |
| M stage (M1) | 2.271 [1.094-4.713] | 0.028^*^ |  | 2.723 [1.249-5.936] | 0.012^*^ |  | 1.672 [0.386-7.247] | 0.492 |  |  |  |
| EGFR^Mem+^ | 0.778 [0.489-1.238] | 0.289 |  |  |  |  | 0.888 [0.341-2.313] | 0.808 |  |  |  |
| EGFR^Cyto+^ | 0.887 [0.575-1.369] | 0.590 |  |  |  |  | 0.460 [0.190-1.112] | 0.085 |  |  |  |
| GPC3^Mem+^ | 1.055 [0.691-1.613] | 0.804 |  |  |  |  | 0.748 [0.309-1.803] | 0.516 |  |  |  |
| GPC3^Cyto+^ | 0.930 [0.610-1.417] | 0.735 |  |  |  |  | 0.646 [0.258-1.620] | 0.352 |  |  |  |
| MUC13^Mem+^ | 0.386 [0.232-0.644] | <0.001^*^ |  | 0.336 [0.050-2.269] | 0.263 |  | 0.641 [0.246-1.672] | 0.363 |  |  |  |
| MUC13^Cyto+^ | 0.446 [0.265-0.751] | 0.002^*^ |  | 1.091 [0.156-7.610] | 0.930 |  | 0.606 [0.220-1.672] | 0.334 |  |  |  |
| ROBO1^Mem+^ | 1.496 [0.980-2.282] | 0.062 |  |  |  |  | 1.649 [0.681-3.996] | 0.270 |  |  |  |
| ROBO1^Cyto+^ | 1.778 [1.077-2.935] | 0.025^*^ |  | 2.367 [1.381-4.055] | 0.002^*^ |  | 1.551 [0.562-4.281] | 0.397 |  |  |  |
| MET^Mem+^ | 1.727 [1.076-2.773] | 0.024^*^ |  | 1.664 [0.951-2.909] | 0.074 |  | 2.335 [0.928-5.876] | 0.072 |  |  |  |
| MET^Cyto+^ | 1.972 [1.277-3.047] | 0.002^*^ |  | 1.095 [0.649-1.847] | 0.733 |  | 2.621 [1.071-6.413] | 0.035^*^ |  | 2.282 [0.880-5.918] | 0.090 |
| CD147^Mem+^ | 1.869 [1.199-2.912] | 0.006^*^ |  | 0.977 [0.587-1.627] | 0.929 |  | 1.588 [0.633-3.983] | 0.325 |  |  |  |
| CD147^Cyto+^ | 2.128 [1.342-3.374] | 0.001^*^ |  | 2.036 [1.168-3.547] | 0.012^*^ |  | 3.430 [1.145-10.275] | 0.028^*^ |  | 3.457 [1.097-10.899] | 0.034^*^ |

HR, Hazard ratio; CI, confidence interval: HBsAg, Hepatitis B surface antigen

^*^Statistically significant (*p*<0.05)

**Table S2.** List of antibodies for flow cytometry

| **Target** | **Isotype** | **Conjugate** | **Clone#/name** | **Vendor** |
| --- | --- | --- | --- | --- |
| CD24 | Mouse IgG1κ | APC | SWA11 | Cell Sciences |
| GPC3 | Human IgG1κ | APC | Codrituzumab | Chugai Pharmaceutical |
| MUC13 | Rabbit IgG | Unconjugated | EPR21901 | Abcam |
| ROBO1 | Rabbit IgG | Unconjugated | Polyclonal | Proteintech |
| TSPAN8 | Rat IgG2b | Unconjugated | 458811 | R&D systems |
| MET | Human IgG1κ | PE | Onartuzumab | Genetech |
| CD147 | Mouse IgG2b | APC | OTI4E4 | OriGene |
| EGFR | Human IgG2κ | APC | Panitumumab | Amgen |
| Mouse IgG | Mouse IgG1 | Unconjuagated | CT6 | Invitrogen |
| Rabbit IgG | Rabbit IgG | Unconjuagated | EPR25A | Abcam |
| Human IgG | Human IgG | FITC | Polyclonal | Novus biologicals |
| Mouse IgG | Goat IgG | Alexa Fluor™ 647 | Polyclonal | Invitrogen |
| Rabbit IgG | Goat IgG | Alexa Fluor™ 647 | Polyclonal | Invitrogen |
| Rat IgG | Goat IgG | APC | Polyclonal | R&D systems |

**Table S3.** Antibodies used for immunohistochemistry

| **Antibody** | **Vendor** | **Clone#** | **Cat. #** | **Incubation** | **Dilution** | **Antigen retrieval** | **Tissue** |
| --- | --- | --- | --- | --- | --- | --- | --- |
| CD24 | Cell Science | SWA11 | FHD65210B | 30 min at RT | 1/2000 | 20 min, PC, pH 6 | Human TMA |
| GPC3 | Cell Marque | 1G12 | 261M-96 | 60 min at RT | 1/500 | 20 min, PC, pH 6 | Human TMA |
| MUC13 | Cell Signaling | E6Z1K | 44454 | 60 min at RT | 1/200 | 20 min, PC, pH 6 | Human TMA |
| ROBO1 | Proteintech | Monoclonal | 67922-1-Ig | 30 min at RT | 1/2000 | 20 min, PC, pH 9 | Human TMA |
| TSPAN8 | Abcam | Polyclonal | ab7007 | 30 min at RT | 1/5000 | 20 min, PC, pH 6 | Human TMA |
| MET | Cell Signaling | D1C2 | 8198 | 60 min at RT | 1/500 | 20 min, PC, pH 9 | Human TMA |
| CD147 | Proteintech | Monoclonal | 66443-1-Ig | 60 min at RT | 1/250 | 20 min, PC, pH 6 | Human TMA |
| EGFR | Dako | H11 | M356301-2 | 60 min at RT | 1/150 | Pro-K, 5 min at RT | Human TMA |
| CD24 | Invitrogen | SN3 | MA5-11828 | 30 min at RT | 1/200 | 20 min, PC, pH 6 | Xenograft |
| GPC3 | Abcam | SP86 | ab95363 | 30 min at RT | 1/3000 | 20 min, PC, pH 6 | Xenograft |
| TSPAN8 | Abcam | Polyclonal | ab7007 | 30 min at RT | 1/1000 | 20 min, PC, pH 6 | Xenograft |
| MET | Cell Signaling | D1C2 | 8198 | 60 min at RT | 1/200 | 20 min, PC, pH 9 | Xenograft |
| CD147 | Thermo Fisher Scientific | 125 | MA5-29060 | 30 min at RT | 1/5000 | 20 min, PC, pH 6 | Xenograft |
| EGFR | Cell Signaling | D38B1 | 4276 | 60 min at RT | 1/100 | 20 min, PC, pH 9 | Xenograft |
| Ki-67 | Abcam | SP6 | Ab16667 | 60 min at RT | 1/200 | 20 min, PC, pH 6 | Xenograft |

RT, room temperature; PC, pressure cooker; Pro-K, proteinase-K
